# Metabolic cooperation among commensal bacteria supports *Drosophila* juvenile growth under nutritional stress

**DOI:** 10.1101/2020.05.27.119370

**Authors:** Jessika Consuegra, Théodore Grenier, Houssam Akherraz, Isabelle Rahioui, Hugo Gervais, Pedro da Silva, François Leulier

## Abstract

The gut microbiota shapes animal growth trajectory in stressful nutritional environments, but the molecular mechanisms behind such physiological benefits remain poorly understood. The gut microbiota is mostly composed of bacteria, which construct metabolic networks among themselves and with the host. Until now, how the metabolic activities of the microbiota contribute to host juvenile growth remains unknown. Here, using *Drosophila* as a host model, we report that two of its major bacterial partners, *Lactobacillus plantarum* and *Acetobacter pomorum* engage in a beneficial metabolic dialogue that boosts host juvenile growth despite nutritional stress. We pinpoint that lactate, produced by *L. plantarum*, is utilized by *A. pomorum* as an additional carbon source, and *A. pomorum* provides essential amino-acids and vitamins to *L. plantarum*. Such bacterial cross-feeding provisions a set of anabolic metabolites to the host, which may foster host systemic growth despite poor nutrition.

**GRAPHICAL ABSTRACT:** 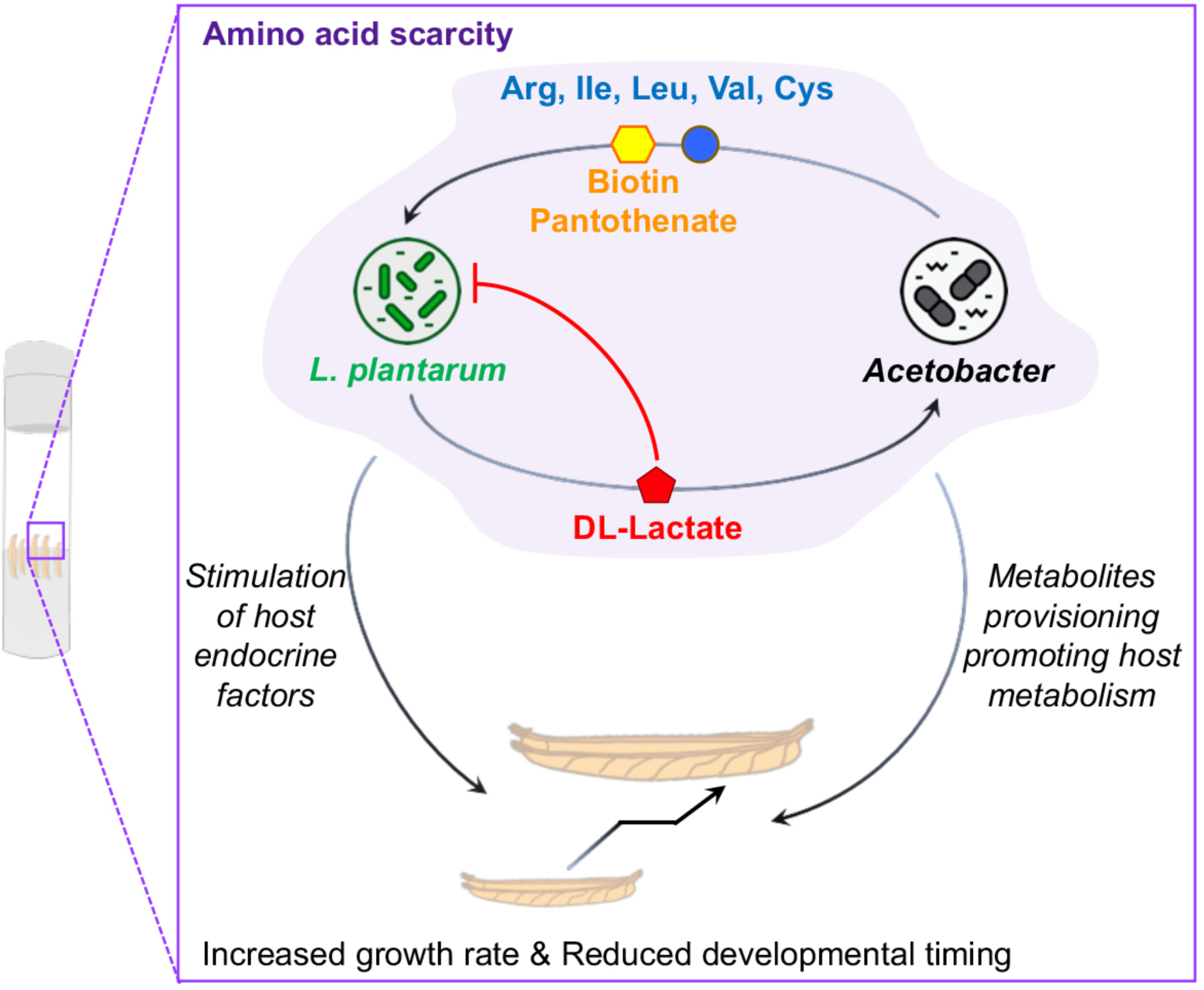

**HIGHLIGHTS:**

- *L. plantarum* feeds lactate to *A. pomorum*
- *A. pomorum* supplies essential amino acids and vitamins to *L. plantarum*
- Microbiota metabolic dialogue boosts Drosophila’s larval growth
- Lactate utilization by *Acetobacter* releases anabolic metabolites to larvae

## INTRODUCTION

In the animal kingdom, juvenile growth takes place during the post-natal stages preceding sexual maturation and ushers in the most profound physiological changes in an organism’s lifetime. These changes are governed by the complex interplay between the animal’s genotype and its nutritional environment. In humans, chronic undernutrition at the juvenile stage leads to severe stunting and long-term negative neurological, metabolic and reproductive consequences (Goyal et al., 2015). Today 155 million children are plagued by childhood malnutrition worldwide (Global Nutrition report, 2018).

Recent studies establish that the microbial communities colonizing the body surfaces (i.e. microbiota), especially the activities and constituents of the gut microbiota, can alter the host’s growth trajectory. Both in invertebrates and in mammals, selected strains of microbiota members can buffer the deleterious impact of undernutrition on juvenile growth dynamics (Blanton et al., 2016; Schwarzer et al., 2016; Shin et al., 2011; Smith et al., 2013; Storelli et al., 2011). In humans, children suffering from malnutrition carry an “immature” gut microbiota that fails to be remedied by classical re-nutrition strategies (Subramanian et al., 2014).

Juvenile growth is marked by the exponential increase of the animals’ biomass manifested as gain in weight and longitudinal size. These physical traits are governed by the host’s growth hormone and growth factors (GH/IGF1 in mammals) whose production and activities are regulated by nutrients availability (Thissen et al., 1994). Recently, it was established that gut microbiota members also influence the production and activity of growth hormone and growth factors in both invertebrate and mammals (Schwarzer et al., 2016; Shin et al., 2011; Storelli et al., 2011; Yan et al., 2016).

Despite recent progress, how the gut microbiota confers such benefits to the host remains poorly understood. This is partly due to the fact that the gut microbiota is a complex ecosystem comprising up to hundreds microbial species in mammals, mostly bacteria (Hooper and Gordon, 2018). They construct multiplex, high-order nutritional and metabolic networks amongst themselves and with the host such that these interactions directly influence host nutrition and metabolism (Schroeder and Bäckhed, 2016). Given this complexity, until now no study has elucidated to what extent and how the metabolic interactions among members of the microbiota contribute to host juvenile growth.

To answer this question, we bypassed the complexity encountered in mammals and developed an experimentally tractable gnotobiotic *Drosophila* model associated with its two major bacterial partners, *Lactobacillus plantarum* and *Acetobacter pomorum*, which are frequently found to co-exist in wild flies captured on fruit-based baits (Chandler et al., 2011; Pais et al., 2018; Wong et al., 2013). Previously, using oligidic diets (i.e., a diet composed of complex ingredients such as inactivated yeast and cornmeal flour), we and others have established that association of germ-free (GF) larvae with either *A. pomorum* or *L. plantarum* stimulates juvenile growth by promoting the systemic release and activities of *Drosophila* Insulin-like peptides (dILPs), the functional analogues of vertebrate Insulin and IGFs (Shin et al., 2011; Storelli et al., 2011). Here, using *Drosophila* bi-associated with *A. pomorum* and *L. plantarum*, we characterized the metabolic dialogues among the three partners in a strictly controlled nutritional environment low in amino acids to mimic chronic protein undernutrition, namely a fully chemically-defined or holidic diet (HD) (Piper et al., 2017). HDs support suboptimal growth and development of *Drosophila* larvae (Jang and Lee, 2018; Piper et al., 2013; Rapport et al., 1983; Schultz et al., 1946) yet it has proved to be a useful tool to study the specific influence of individual nutrients on *Drosophila* physiology (Jang and Lee, 2018; Mishra et al., 2018; Piper et al., 2013, 2017). This experimental model grants us complete control over 3 key parameters in the system: the diet, the host and its commensal partners. We defined the nutritional requirements, auxotrophies and complementation of over 40 individual nutrients including all amino acids, vitamins, nucleic acids, lipid precursors and minerals for each commensal and the juvenile host in the GF context or upon association with either microbial partner (Consuegra et al., 2020).

Here, we report that when co-inoculated on a *Drosophila* HD low in amino acids, *L. plantarum* and *A. pomorum* engage in a beneficial metabolic dialogue that supports bacterial growth and buffers the deleterious impact of nutritional stress on host juvenile growth. We specifically pinpoint that lactate, the main metabolic by-product of *L. plantarum*, is utilized by *A. pomorum* as an additional carbon source, and in turn, *A. pomorum* provides various amino-acids and B-vitamins to complement *L. plantarum* auxotrophies. Inert microbial biomass has been reported to promote larval development (Bing et al., 2018; Storelli et al., 2011) and adult longevity (Keebaugh et al., 2018; Yamada et al., 2015) probably by acting as an additional nutritional source. While we confirm that inert bacterial biomass slightly contributes to increased juvenile growth, we show that *Lactobacillus* provision of lactate to *Acetobacter* triggers a metabolic shift in *Acetobacter* leading to the provision of a set of anabolic metabolites to the host, which may boost host systemic growth despite poor nutrition.

## RESULTS

### Bi-association enhances the benefit of commensal bacteria on larval development

In a Holidic Diet (HD) low in amino acids that mimics chronic protein undernutrition, we studied larval development in germ-free (GF) and upon mono or bi-association with two representative commensal strains of the *Drosophila* microbiota: *Acetobacter pomorum*^WJL^ (Ap^WJL^) and *Lactobacillus plantarum*^NC8^ (Lp^NC8^). In this diet, GF larvae reach metamorphosis at ~10 days. By comparison, the time from embryogenesis to metamorphosis of GF animals on rich oligidic diets (i.e. yeast, 50 g/L) is ~5 days while it is increased to ~13 days on poor oligidic diet (i.e. yeast, 6g/L) (Matos et al., 2017).

On HD, the benefit on larval development of bacterial mono-association is enhanced in larvae bi-associated with Ap^WJL^ and Lp^NC8^ (Ap^WJL^:Lp^NC8^; Fig. 1A-B). Bi-associated animals always develop faster than their mono-associated siblings and reach metamorphosis in ~5.2 days (Fig. 1A) or ~8.2 day (Fig. 1B) according to the initial bacterial inoculum. We observed similar results using both complete HDs with optimal amino acid content (Fig. S1A, HD 16g and HD 20g) or with a fruit-based diet (Banana diet, Fig. S1B) containing ~7 g/kg of protein (Oyeyinka and Afolayan, 2019) where GF larvae fail to develop (see Methods). Of note, the differential capacities of the bacteria to sustain *Drosophila* growth on the banana diet are not a consequence of differential bacteria growth on this fruit-based diet as both Ap^WJL^ and Lp^NC8^ grew to the same extent in presence or absence of larvae (Fig. S1C-D).

**Fig. 1:**
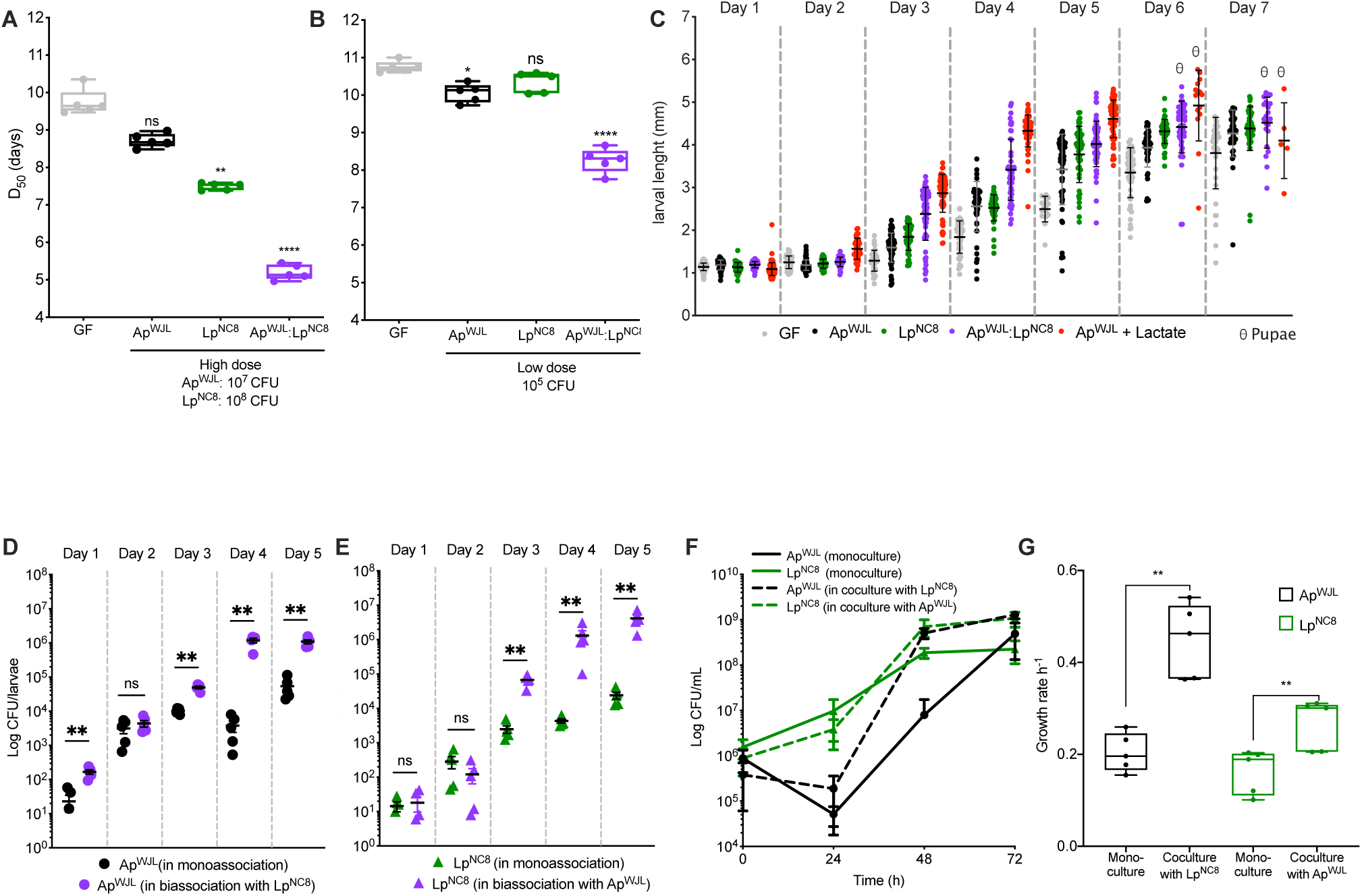
Bi-association with Ap^WJL^ and Lp^NC8^ enhances commensal mediated benefit on larval development. (A-B) Developmental timing (time from egg to metamorphosis) on complete holidic diet (HD) of Germ Free (GF) larvae (grey) or GF larvae inoculated with high dose (10^7^ or 10^8^ CFU respectively; A) or low dose (10^5^ CFU; B) of Ap^WJL^ and/or Lp^NC8^ (Ap^WJL^ – black, Lp^NC8^ – green/Ap^WJL^:Lp^NC8^ – purple). D_50_: Day when 50% of the larvae population has entered metamorphosis. (C) Larval length at every day post-embryogenesis of GF larvae or post-inoculation (Day 1) with 10^5^ CFU of Ap^WJL^ and/or Lp^NC8^ or Ap^WJL^ mono-associated larvae supplemented with DL-lactate at a final concentration of 0.6 g/L (red). θ: pupae detected in the vial. (D-E) Microbial load (Ap^WJL^, D; Lp^NC8^, E) of larvae mono- or bi-associated with 10^5^ CFU of Ap^WJL^ and/or Lp^NC8^. (F-G) Growth in liquid HD (F) and growth rates (G) of Ap^WJL^ and Lp^NC8^ in mono-(plain lines) or co-cultures (dashed lines) in liquid HD. Grey always refers to GF, black to Ap^WJL^ mono-association, green Lp^NC8^ mono-association condition and purple Ap^WJL^:Lp^NC8^ bi-association. Each symbol represents an independent replicate apart in (F) where symbols represent the means ± SEM of three biological replicates. Boxplots show minimum, maximum and median where each point is a biological replicate. Dot plots show mean ± SEM. (A-B) We performed Kruskal-Wallis test followed by uncorrected Dunn’s tests to compare each gnotobiotic condition to GF. (D-E) Each point represents a biological replicate comprising the average microbial load of a pool of 10 larvae. We performed Mann-Whitney test to compare microbial loads in mono-association to microbial loads in bi-association for the strain of interest at each time point. (G) We performed Mann-Whitney test to compare the growth rate in monoculture to the growth rate in coculture for the strain of interest. ns: non-significant, *: p-value<0,05, **: p-value<0,005, ***: p-value<0,0005, ****: p-value<0,0001. See also Figure S1 and S2.

During post-embryonic development, Ap^WJL^ or Lp^NC8^ not only influences maturation rates (i.e. time to entry to metamorphosis), but also increases larval linear size gains upon nutrient scarcity (Fig. 1C). Ap^WJL^:Lp^NC8^ bi-association also enhances the benefit of commensals on this trait as early as 3 days after bi-association (Fig. 1C).

Next, we wondered if each bacterium benefits from the presence of the other. To this end, we assessed the microbial load in larvae through larval development upon mono- and bi-association with Ap^WJL^, Lp^NC8^ or Ap^WJL^:Lp^NC8^, respectively. Ap^WJL^ and Lp^NC8^ loads in mono- or bi-association start to differ from day 3 after egg laying and reach a two-log difference at day 5 (Fig. 1D, E). The reciprocal benefit between Ap^WJL^ and Lp^NC8^ is also observed while bacteria grow in a liquid version of the HD (see Methods). In co-culture, Ap^WJL^ and Lp^NC8^ have slightly higher final biomasses (Fig. 1F) and marked higher growth rates (Fig. 1G) than in mono-cultures. As previously reported in other experimental settings, the enhanced benefit of commensals on fly’s lifespan (Yamada et al., 2015) or larval development (Bing et al., 2018; Storelli et al., 2011) is mediated at least partly by the trophic effect of providing inert microbial biomass as nutrients to the host. Since we detected a slight increased bacterial biomass in the diet and the host upon bi-association, we investigated the contribution of such inert biomass to the observed growth promotion phenotype. To this end, we inoculated GF larvae with Heat Killed (HK) Ap^WJL^ or Lp^NC8^ at high dose (10^9^ CFU) in mono- and bi-associated conditions (Fig. S2A). Mono-association with HK bacteria at high or low doses fails to accelerate larval development (Fig. S2A, B), yet bi-association with HK bacteria at high doses slightly contribute to host development by accelerating larval development by ~1 day compared to GF animals (Fig. S2A). However, this effect is very mild when compared to the effect of live and metabolic active bacteria bi-association at high or low doses (Fig. 1A, B), which respectively led to larval development accelerations of ~5.5 or ~2.5 days compared to GF conditions. Of note, in contrast to live bacteria bi-association, bi-association with HK bacteria on HDs with an increased amino acids content or a banana diet did not rescue or accelerate larval development (Fig. S1A-B). Moreover, the enhanced *Drosophila* growth observed upon bi-association requires both bacteria to be metabolically active and associated to the host from early stages of development, since bi-association where one of the bacteria is HK (Fig. S2B) or delayed bi-association (Fig. S2C, D) fails to accelerate larvae development.

Collectively, our results show that microbial bi-association of larvae developing in a suboptimal nutritional context results in increased host’s maturation rates and size gains compared to mono-associations. This beneficial effect partially results from a trophic effect of increase bacterial biomass provision to the host but mostly relies on the functional impact of alive and metabolically active microbes.

### Ap^WJL^ benefits Lp^NC8^ via essential amino acid and vitamins provision

Recently, we showed that Ap^WJL^ and Lp^NC8^ differentially fulfills the nutritional requirements of the ex-GF larva thanks to their individual genetic repertoires. In this context, the positive impact of Ap^WJL^ or Lp^NC8^ on host development requires metabolically active bacteria, and is independent of bacterial loads in the depleted diets or in the larval gut (Consuegra et al., 2020). Specifically, we identified the nutritional auxotrophies of both Ap^WJL^ and Lp^NC8^ in HD. Ap^WJL^ is completely prototroph, whereas Lp^NC8^ is auxotroph for Arg, Ile, Leu, Val, Cys, Biotin and Pantothenate. Such differences between Ap^WJL^ and Lp^NC8^ were expected. Indeed, *L. plantarum* is a fastidious bacterium with complex metabolic requirements including amino acids and vitamins (Martino et al., 2016; Vos et al., 2009). Therefore, in a simple microbial community like the one studied here, a prototrophic bacterium like *A. pomorum* may support *L. plantarum* growth by providing essential amino acids and vitamins.

To directly test this hypothesis, we studied the growth of Lp^NC8^ in the presence of Ap^WJL^ in liquid HD lacking each of the amino acids and vitamins for which it is auxotroph. We set monocultures of Ap^WJL^ and Lp^NC8^ and a coculture of Ap^WJL^:Lp^NC8^ in liquid HDΔArg, ΔIle, ΔLeu, ΔVal, ΔCys, ΔBiotin or ΔPantothenate and assessed the bacterial counts in mono and cocultures during 72h. As expected, Ap^WJL^ grows in these media to the same extent as in the complete HD, whereasLp^NC8^ is unable to grow as a monoculture (Fig. 2A-G). Interestingly, Lp^NC8^ grows in the deficient media only when cocultured with Ap^WJL^ (Fig. 2A-G). From the HDΔArg, HDΔIle and HDΔLeu mono- and cocultures, we also recovered supernatants and quantified Arg, Ile and Leu release in the media using High-Performance Liquid Chromatography (HPLC). In Ap^WJL^ monocultures, we observe an accumulation of these amino acids that correlates with Ap^WJL^ growth (Fig. 2H-J). As expected, they are not detected in the Lp^NC8^ monocultures (Fig. 2H-J). In Ap^WJL^:Lp^NC8^ coculture, we do not detect any accumulation of Arg or Leu and a reduction in Ile accumulation, which suggests that the amino acids released by Ap^WJL^ are immediately consumed by Lp^NC8^ to support its growth and thus do not accumulate in the media (Fig. 2H-J). These results therefore establish that Ap^WJL^ provides amino acids, and probably B vitamins to Lp^NC8^.

**Fig. 2:**
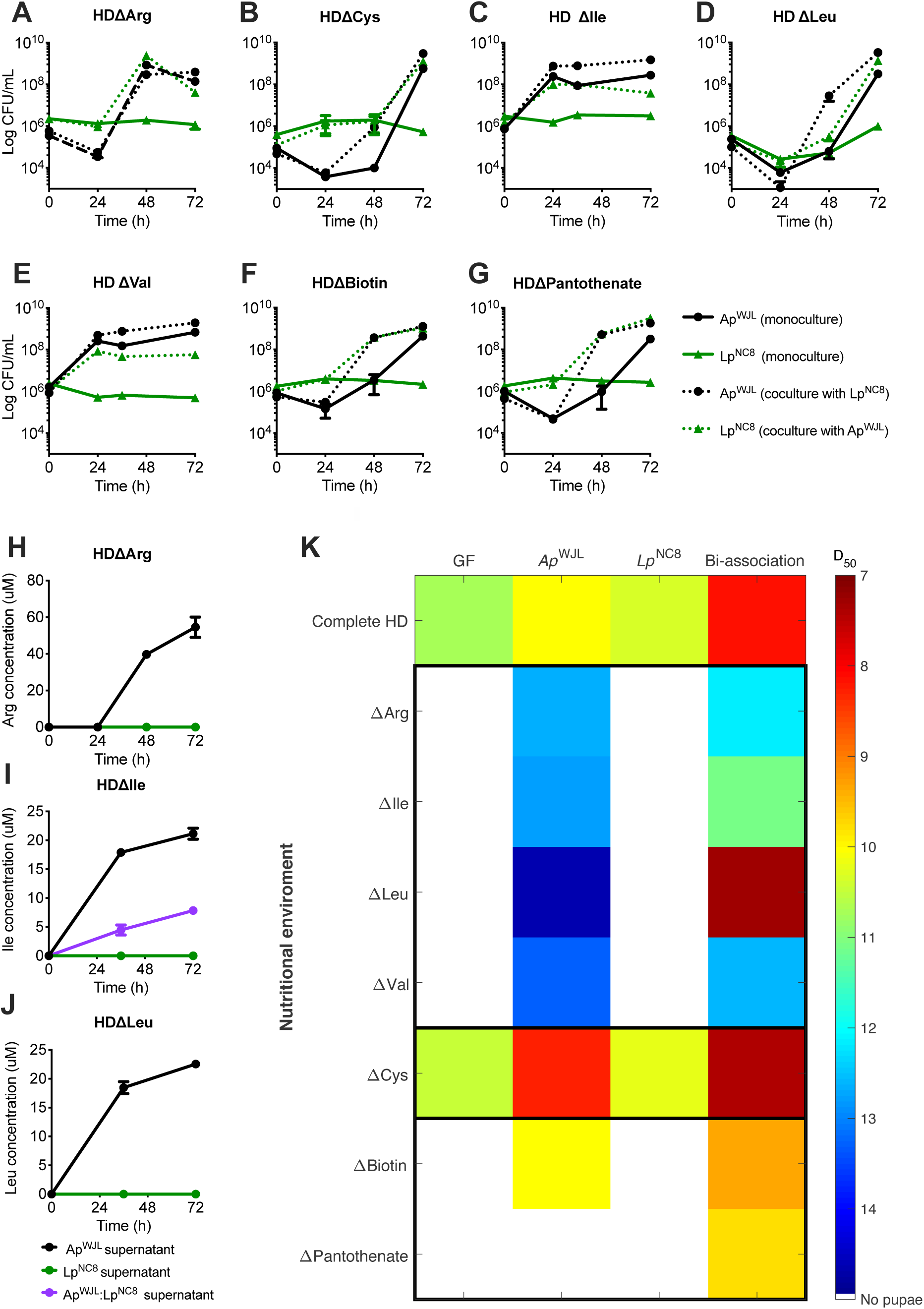
Ap^WJL^ benefits Lp^NC8^ via essential amino acid and vitamins provision. (A-G) Growth curves of Ap^WJL^ and Lp^NC8^ in mono-(plain lines) or co-cultures (dotted lines) in liquid holidic diets (HD) lacking either Arg (HDΔArg, A), Cys (HDΔCys, B), Ile (HDΔIle, C), Leu (HDΔLeu, D), Val (HDΔVal, E), Biotin (HDΔBiotin, F) or Pantothenate (HDΔPantothenate, G). Black refers to Ap^WJL^, green the Lp^NC8^. (H-J) HPLC quantification of Arg, Ile and Leu in Ap^WJL^ or Lp^NC8^ mono-culture supernatants (blak and green lines, respectively) or Ap^WJL^:Lp^NC8^ co-culture (purple line) in HDΔArg, HDΔIle, HDΔLeu, respectively. (A-J) Symbols represent the means ± SEM of three biological replicates. (K) Heat map representing the mean D_50_ (day when 50% of the larvae population has entered metamorphosis) of GF larvae (first column) and larvae mono-associated with Ap^WJL^ or Lp^NC8^ or bi-associated with Ap^WJL^:Lp^NC8^ (columns 2, 3 and 4 respectively). Each row shows D_50_ in a different version of the HD: complete HD or HDs each lacking a specific nutrient HDΔArg, HDΔIle, HDΔLeu, HDΔVal, HDΔCys, HDΔBiotin, HDΔPantothenate. White color code means that larvae did not reach pupariation.

### Ap^WJL^ to Lp^NC8^ nutrient provision potentiate commensal mediated larval auxotrophies compensation

Next, we sought to determine if these metabolic interactions among *Drosophila* commensals could be translated into a further benefit to larvae developing on media lacking each of the amino acids and vitamins for which Lp^NC8^ is auxotrophic. We therefore assessed the developmental time in HDArg, Ile, Leu, Val, Cys, Biotin and Pantothenate of mono-(Ap^WJL^ or Lp^NC8^) or bi-associated (Ap^WJL^:Lp^NC8^) larvae (Fig. 2K). Association of the larval host with Ap^WJL^ compensates all nutrient depletions except for pantothenate, while Lp^NC8^ fails to compensate the lack of any nutrient for the host due to its own auxotrophies. Interestingly, bi-association with Ap^WJL^:Lp^NC8^ systematically exceeds the benefit provided to the host by mono-association with Ap^WJL^, and in HDΔPantothenate even rescues host viability (Fig. 2K).

Taken together, these results establish that upon bi-association, Ap^WJL^ supplies Arg, Ile, Leu, Val, Cys, biotin and pantothenate to Lp^NC8^, thus allowing both commensals to thrive on these depleted media. This nutritional cooperation then potentiates the commensal-mediated promotion of larval development in depleted diets via the bacterial provision of the missing essential nutrients to the host.

### Lp^NC8^ derived lactate benefits Ap^WJL^ and enhances Ap^WJL^ mediated larval growth promotion

Next, we wondered how Ap^WJL^ benefits from Lp^NC8^ (Fig. 1F, G). We hypothesize that Lp^NC8^ metabolic by-products enhance the ability of Ap^WJL^ to promote larval development. To test this, we mono-associated GF embryos with Ap^WJL^ and added either sterile PBS or the supernatant of a culture of Lp^NC8^ grown on liquid HD for 3 days. The addition of a Lp^NC8^ supernatant on embryos mono-associated with Ap^WJL^ is sufficient to accelerate larval development by ~4 days compared to GF animals, while Ap^WJL^ mono-association only triggers a single day acceleration. However, addition of Lp^NC8^ supernatant did not improve larval development in GF condition or in mono-association with Lp^NC8^ (Fig. 3A).

**Fig. 3:**
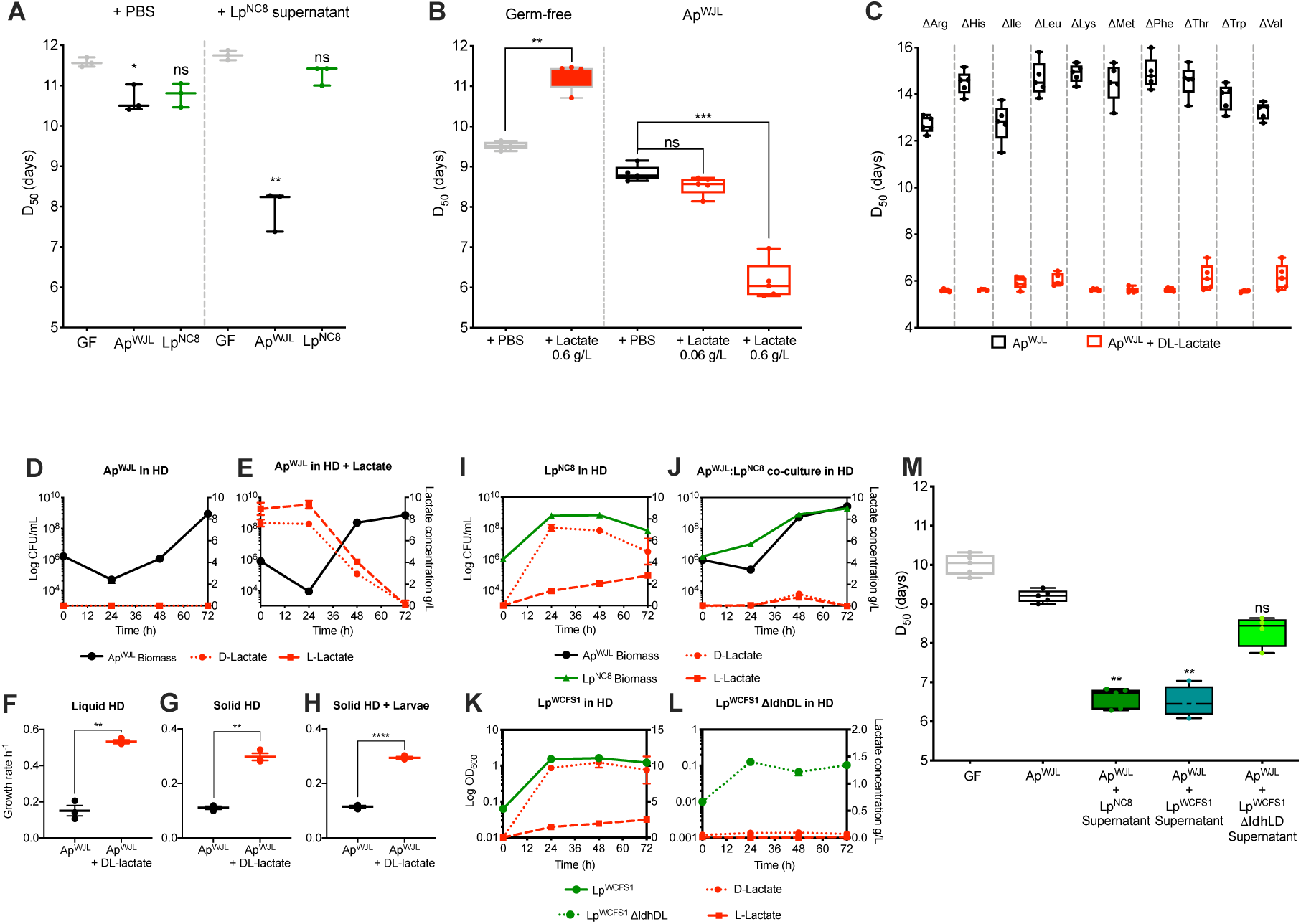
Lp^NC8^ derived lactate benefits Ap^WJL^ and enhance Ap^WJL^ mediated larval growth promotion. (A, M) Developmental timing of Germ Free (GF) larvae or GF larvae inoculated with 10^5^ CFU of Ap^WJL^ (A, M black) or Lp^NC8^ (A, green) supplemented with either sterile PBS (A) or the supernatant from a 72h culture of Lp^NC8^ (A, M), LpWCSF1 (M, turquoise) or Lp^WCFS1^ΔldhDL (M, light green) in complete holidic diet (HD). (B) Developmental timing on HD of GF larvae (grey) or GF larvae inoculated with 10^5^ CFU of Ap^WJL^ supplemented with either sterile PBS (black) or DL-lactate solutions (red) at inoculation (final concentration in the diet 0.06 g/L or 0.6 g/L). (A, B, M) Each dot represents an independent biological replicate. Boxplots show minimum, maximum and median. We performed Kruskal-Wallis test followed by uncorrected Dunn’s tests to compare each condition to GF. ns: non-significant, *: p-value<0,05, **: p-value<0,005, ***: p-value<0,0005. (C) Developmental timing of GF larvae inoculated with 10^5^ CFU of Ap^WJL^ supplemented at inoculation with either sterile PBS (black) or DL-lactate at final concentration of 0.6 g/L in HDs lacking each an essential amino acid for *Drosophila*: from left to right, HDΔArg, HDΔHis, HDΔIle, HDΔLeu, HDΔLys, HDΔMet, HDΔPhe, HDΔThr and HDΔVal. Boxplots show minimum, maximum and median and each dot represents an independent biological replicate. Growth curves (D-E), and growth rates (F) of Ap^WJL^ in liquid HD supplemented (E) or not (D) with DL-lactate solution. D-(dotted line) and L-lactate (dashed line) levels (red) were quantified in both conditions. Growth rates of Ap^WJL^ in solid HD and HD + DL-lactate with (H) or without (G) larvae. (I-L) Growth curves in liquid HD of Lp^NC8^ (green) or Ap^WJL^ (black) in mono-(I) or co-culture (J), or LpWCSF1 (K, green) or Lp^WCFS1^ΔldhDL (L, dotted green) with the respective D-(dotted line) or L-lactate (dashed line) levels (red). Note the low OD_600_ of Lp^WCFS1^ΔldhDL vs Lp^WCSF1^ but similar CFU counts (Fig. S4A-B). Symbols represent the means ± SEM of three biological replicates except for panel (F-H) where each symbol represents an independent replicate ± SEM. See also Figure S1, S3 and S4.

*L. plantarum* is a homolactic fermentative microorganism that secretes its principal metabolic by-products: D- and L-lactate into the nutritional substrate. We next assayed if an equimolar solution of DL-lactate could reproduce the benefit of Lp^NC8^ supernatant on embryos mono-associated with Ap^WJL^. When DL-lactate is added at a final concentration of 0.6 g/L, larvae mono-associated with Ap^WJL^ exhibit strong developmental acceleration and linear size gain (Fig. 3B and 1C). However, DL-lactate is deleterious to GF larvae as it delays development by ~2 days (Fig. 3B). Furthermore, in HD lacking each of the fly essential amino acids (Fig. 3C) or in complete HDs with optimal amino acid content (Fig. S1A, HD 16g and HD 20g), the DL-lactate supplementation to larvae mono-associated with Ap^WJL^ reproduces and even exceeds the benefit of the bi-association.

*A. pomorum* is an acetic acid bacterium that produces acetic acid by aerobic fermentation. We first confirmed that Ap^WJL^ does not produce lactate during growth on liquid HD (Fig. 3D) but is capable of consuming exogenous sources of lactate in the cultured media, without a preference of either chiral form (Fig. 3E). Consumption of DL-lactate by Ap^WJL^ slightly increases its final biomass in solid HD (Fig. S3A-B), reaching an average ~4×10^7^ CFU/tube (instead of ~1×10^7^ CFU/tube when lactate was omitted) and markedly enhances bacterial growth rate in both liquid (Fig. 3F) and solid HD with or without larvae (Fig. 3G-H, Fig. S3A-B). In liquid HD, we quantified that Lp^NC8^ releases ~8 g/L of DL-lactate (3:1 ratio, D:L; Fig. 3I). Finally, in an Ap^WJL^:Lp^NC8^ coculture, we observed that the lactate released by Lp^NC8^ is immediately consumed by Ap^WJL^, preventing its accumulation in the media (Fig. 3J).

Next, we wondered if the beneficial effect on larval development we observed upon supplementation with DL-lactate of Ap^WJL^ mono-associated larvae is due to the mere increase on Ap^WJL^ biomass. To test this hypothesis, we assessed the development of larvae mono-associated with Ap^WJL^ in two conditions: first, with a high dose of Ap^WJL^ biomass (~10^8^ CFU) so it matches the final bacterial count at stationary phase in solid HD supplemented with lactate in the presence of larvae. Second, live Ap^WJL^ biomass associated to *Drosophila* larvae was corrected daily to match the biomass reached when Ap^WJL^ mono-associated animals are supplemented with lactate, according to the bacteria growth dynamics established in Fig. S3B (Fig. S3C, S3D). Mono-association with a higher dose of Ap^WJL^ (10^8^ CFU) was deleterious to larval development (Fig. S3D), this also justifies our choice of 10^7^ CFU Ap^WJL^ inoculum in Fig.1A. Indeed, in two out five replicates, flies did not reach pupariation (egg-to-pupae survival <20%, Fig. S3E). In the other 3 replicates, egg-to-pupae survival was higher (~80%) as well as variability among replicates (CV=17.4%). In the Ap^WJL^ lactate-matched biomass condition, larval development was not faster than larvae mono-associated with Ap^WJL^, yet lactate supplementation triggered the expected enhanced larval development of Ap^WJL^ mono-associated animals (Fig. S3D). Thus, we conclude that the enhanced host growth observed upon lactate supplementation to Ap^WJL^ is not due to the mere increase in Ap^WJL^ biomass and growth rate upon lactate consumption.

The lactate produced by Lp^NC8^ seems to be the key metabolite altering Ap^WJL^ metabolism and its influence on host growth. To directly test this hypothesis, we recovered supernatants of 3-days cultures in liquid HD of a *L. plantarum* strain lacking the *ldh* genes (Lp^WCFS1^Δl*dhDL*) and its wild-type counterpart (Lp^WCFS1^) and assessed their effects on the development of larvae mono-associated with Ap^WJL^. Lp^WCFS1^Δl*dhDL* has been reported to produce only traces amounts of D- and L-lactate (Ferain et al., 1996). We confirmed these findings in liquid HD by monitoring bacterial growth and DL-lactate production by both strains for 72 hours (Fig. 3K, L). Both strains grow in MRS and liquid HD to the same extent without any difference in their final biomass (CFU/mL) despite the observed reduced OD_600_ of Lp^WCFS1^Δl*dhDL* (Fig. S4A-B). Lp^WCFS1^ supernatant at 72h contains ~9.4 g/L of D-lactate and ~2.5 g/L of L-lactate (Fig. 3K). Lp^WCFS1^Δl*dhDL*, on the other hand, only accumulates a total of ~0.09 g/L of DL-lactate (Fig. 3L). Importantly, as in a HD + DL-lactate, Ap^WJL^ growth rate is higher when growing on a Lp^NC8^ or Lp^WCFS1^ supernatants but not on Lp^WCFS1^*ldhDL* supernatant (Fig. S4C). Also, lactate or lactate-containing supernatants from Lp^NC8^ or Lp^WCFS1^ sustain increased Ap^WJL^ larval loads during development (Fig. S4D), as does bi-association with Lp^NC8^ (Fig. 1D). Finally, the addition of a supernatant from Lp^WCFS1^ culture on larvae mono-associated with Ap^WJL^ boosts larval growth and maturation to a degree comparable to Lp^NC8^‘s supernatant (Fig. 3M). The effect of these supernatants on host development is not due to secreted bacterial peptides since the total amino acids’ concentration of Lp^NC8^ culture supernatants remain stable during growth on liquid HD (Fig. S4E), and the addition of an equal volume of sterile liquid HD (containing an amount of amino acids similar to the culture supernatant) on larvae mono-associated with Ap^WJL^ does not accelerate development (Fig. S4F). Instead, the impact of the tested supernatants on larval development is most likely due to the lactate produced by Lp^NC8^ and Lp^WCFS1^ (Fig. 3I, K) since a supernatant from Lp^WCFS1^Δl*dhDL* culture fails to accelerate development of larvae mono-associated with Ap^WJL^ (Fig. 3M).

So far, we demonstrated that the positive effect of *L. plantarum* supernatant on larva mono-associated with Ap^WJL^ is based on its lactate content. Importantly, treatment of GF larvae with either the supernatants of Lp^WCFS1^ or Lp^WCFS1^Δ*ldhDL* has no effect on GF larvae development, neither does treatment with either a supernatant of Ap^WJL^ grown in the presence of these filtrates or with filtrates of Ap^WJL^ cocultured with any of the test *L. plantarum* strains (Fig. S4G). Therefore, we first conclude that DL-lactate does not directly benefit the larval host, rather DL-lactate may trigger a switch of carbon utilization in Ap^WJL^, which in turn reconfigures the metabolic by-products it releases, which the host utilizes to fuel its anabolic growth.

### Lactate mediated enhanced Ap^WJL^ larval growth promotion does not rely on amino acid provision to the host

To test our proposal, we focused on lactate metabolism in *A. pomorum*. Unfortunately, little is known about the core metabolism of this *Acetobacter* species. Most metabolic and genetic studies on *Acetobacter* have been performed on *A. aceti* due to its industrial use on vinegar production (Sakurai et al., 2010) or on *A. pasterianus* as a core member of the fermenting microbiota of cocoa (Adler et al., 2014) which shares ~90% nucleotide identity with *A. pomorum* (Sannino et al., 2018). *A. pasterianus* oxidizes lactate to pyruvate and converts it to (1) acetoin, which is released into the surrounding media, to (2) acetyl-CoA which is directed to the TCA cycle, or (3) to phosphoenolpyruvate (PEP) for gluconeogenesis. In the two last cases, lactate consumption is accompanied by higher metabolic fluxes through biosynthetic pathways for biomass production including *de novo* amino acids biosynthesis (Adler et al., 2014).

We thus wondered if lactate consumption by Ap^WJL^ triggers an increased production and release of amino acids that would be consumed by the host and would stimulate larval growth. To test this hypothesis, we set cultures in liquid HD with or without DL-lactate supplementation, followed bacterial counts and sampled supernatants every 24h for 72 hours for quantification of amino acids. We calculated the net amino acids release in each condition at 24h, 48h and 72h by subtracting the amino acid concentration quantified at 0h from each incremental time points (Fig. 4A, B). First, we observed a distinct release of amino acids at 24h and 48h in both conditions. In the absence of lactate, we focused on the amino acids release by Ap^WJL^ at 48h, while in the middle of its exponential phase (Fig. 4A inner panel). With DL-lactate addition (Fig. 4B), we observed a distinct release of amino acids at 24h (early exponential phase) and 48h (late exponential phase, Fig. 4B inner panel). Unexpectedly, during the stationary phase at 72h, amino acids are depleted instead of accumulation.

**Fig. 4:**
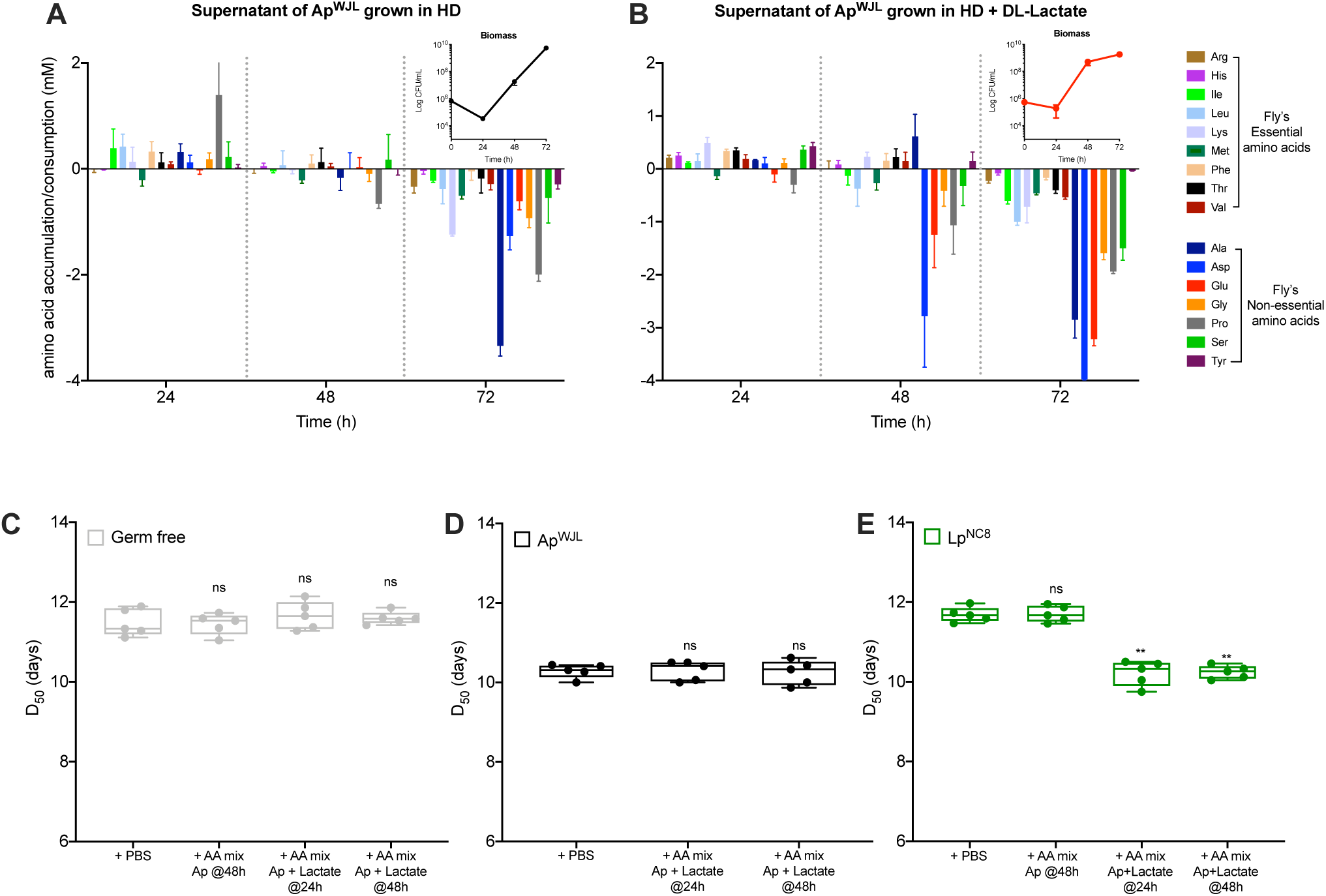
Upon lactate consumption Ap^WJL^ produces an amino acid cocktail that enhances the growth promoting ability of Lp^NC8^. (A-B) Net production of essential and non-essential fly amino acids at 24h, 48h and 72h. Net production was calculated from HPLC quantification data by subtracting the amino acid concentration quantified at 0h from each incremental time points. Conditions included the supernatant of Ap^WJL^ cultures (inner panels) in complete HD supplemented (B) or not (A) with DL-lactate. (C-E) Developmental timing of GF larvae (C) inoculated with 10^5^ CFU of Ap^WJL^ (D) or 10^5^ CFU of Lp^NC8^ (E) supplemented with either sterile PBS, the amino acid mix produced by Ap^WJL^ in liquid culture at 48h (+AA mix Ap @48h), the amino acid mix produced by Ap^WJL^ in liquid culture supplemented with DL-lactate at 24h (+AA mix Ap +Lactate @24h) or the amino acid mix produced by Ap^WJL^ in liquid culture supplemented with DL-lactate at 48h (+AA mix Ap +Lactate @48h). See Table S1 for detailed information on the amino acid mixes. Boxplots show minimum, maximum and median, each point represents a biological replicate. We performed Kruskal-Wallis test followed by uncorrected Dunn’s tests to compare each condition to the PBS treated condition. ns: non-significant, **: p-value<0,005.

Based on these observations, we prepared solid HDs each supplemented with the specific concentration of amino acids mixtures from each specific time points (Table S1; See Methods). These include: a mixture of the amino acids representative of those released by Ap^WJL^ in liquid HD at 48h (AA mix Ap @48h) and the mixtures of the amino acids released by Ap^WJL^ at 24 and 48h in liquid HD supplemented with DL-lactate (AA mix Ap+lactate @24h and AA mix Ap+lactate @48h, respectively) (Fig. 4B and inner panel). We then assessed the maturation time of GF and Ap^WJL^ mono-associated larvae on these three supplemented diets. We observe no enhanced benefit of the different amino acid mixes on GF or Ap^WJL^ mono-associated larvae maturation time (Fig. 4C, D).

These results suggest that amino acids release by Ap^WJL^ is not a key mechanism by which Ap^WJL^ promotes host growth on complete HD, but we cannot rule out the contribution of amino acid precursors or derivatives to host growth promotion in this setting. However, our results indicate that the enhanced beneficial effect of Ap^WJL^ on larval development upon DL-lactate metabolization is not mediated by *de novo* amino acid biosynthesis and release.

### Upon lactate consumption Ap^WJL^ produces amino acids that enhance the growth promoting ability of Lp^NC8^

We previously established that Ap^WJL^ cross-feeds amino acids and B-vitamins to Lp^NC8^ (Fig. 2). Therefore, we wonder if the amino acids mix produced by Ap^WJL^ while growing on HD supplemented with DL-lactate would further enhance the larval growth promotion ability of Lp^NC8^. We tested this hypothesis in the same set-up described above (Fig. 4A-D). We prepared solid HDs supplemented with the three different mixtures of amino acids (AA mix Ap @48h; AA mix Ap+lactate @24h and AA mix Ap+lactate @48h; Table S1). On these three supplemented media, the development of Lp^NC8^ mono-associated larvae is significantly accelerated with either the AA mix Ap+lactate @24h or AA mix Ap+lactate @48h but not with the AA mix Ap @48h (Fig. 4E).

Together our results indicate that upon consumption of the DL-lactate secreted by Lp^NC8^, Ap^WJL^ releases amino acids that are now accessible to Lp^NC8^. As a result, these amino acids further benefit Lp^NC8^ and enhance Lp^NC8^ mediated larval growth promotion in complete HD. However, the amino acids released by Ap^WJL^ in response to lactate do not directly influence the host. This is therefore the metabolic cooperation between the two commensals that results in increased host juvenile growth, higher microbial larval loads (Fig. 1D, E) and improved growth rate of Ap^WJL^ and Lp^NC8^ in the HD (Fig. 1F, G). These results establish that the metabolic cooperation occurring between the two major commensal bacteria of *Drosophila* support an optimal nutritional mutualism among all the partners while facing amino acid scarcity.

### Lactate utilization by *Acetobacter* is necessary to its physiological response to Lp^NC8^ and enhanced benefit on host growth

We aimed to elucidate the mechanisms underpinning the *Lactobacillus*-derived lactate influence on *Acetobacter* in relation to its increased potential to mediate larval growth. First, we focused on lactate utilization by *Acetobacter*. As mentioned previously, DL-lactate consumption by *A. pasterianus* generates acetoin and an increased carbon flux towards gluconeogenic pathways. These metabolic features seem to be shared among other *Acetobacter* species such as *A. fabarum*^DsW_054^ (Af), a strain isolated from wild-caught *Drosophila suzukii* (Winans et al., 2017). Indeed, Sommer & Newell recently reported that lactate produced by *L. brevis* is metabolized by Af through gluconeogenesis pathways via LDH and PPDK, while pyruvate is converted to acetoin by ALS and ALDC (Sommer and Newell, 2018) (Fig. 5A). Based on this information, we hypothesized that the effect of DL-Lactate on Ap^WJL^ and the development of Ap^WJL^ mono-associated larvae relies on the lactate utilization by Ap^WJL^ and its conversion to acetoin or to an increased flux towards gluconeogenic pathways (Fig. 5A). To test these hypotheses, we use a set of Af mutants affecting key enzymes of the lactate metabolism from the Af’s transposon insertion mutant library generated by (White et al., 2018) (Fig. 5A). First, we confirmed that in HD Af behaves like Ap^WJL^. As Ap^WJL^, Af tends to accelerate larval development and Lp^NC8^ supernatant or DL-lactate supplementation all enhance the influence of Af on larval growth (Fig. 5B-C). As Ap^WJL^, Af also consumes exogenous sources of DL-lactate, without a preference of either chiral form (Fig. S5A). Af prevents accumulation of DL-lactate produced by Lp^NC8^ when cocultured with this strain in liquid HD (Fig. S5B). The first step of lactate metabolism is its oxidation by the enzyme lactate dehydrogenase (LDH) to produce two H+ and pyruvate (Fig. 5A). We tested two independent Af mutants in the *ldh* gene (Af::Tn*ldh*, clones 10B7 and 92G1; (Sommer and Newell, 2018; White et al., 2018). These mutants grow in liquid HD to the same extent than the Af wild-type strain (Fig. S5C). On a HD supplemented with DL-lactate, Af::Tn*ldh* mutants consume the D-chiral form of lactate (D-Lactate) (Fig. S5D-E) and still confer a significant benefit to larvae development upon addition of either DL-lactate or D-lactate, albeit with a slight reduction as compared to the WT strain (Fig. 5C). However, both Af::Tn*ldh* mutants fail to consume L-lactate (Fig. S5D-E) and accordingly completely fail to enhance larvae development upon addition of L-lactate (Fig. 5C). These results therefore stablish that the positive effect of lactate on the development of *Acetobacter* mono-associated larvae relies on lactate utilization by *Acetobacter* strains.

**Fig. 5:**
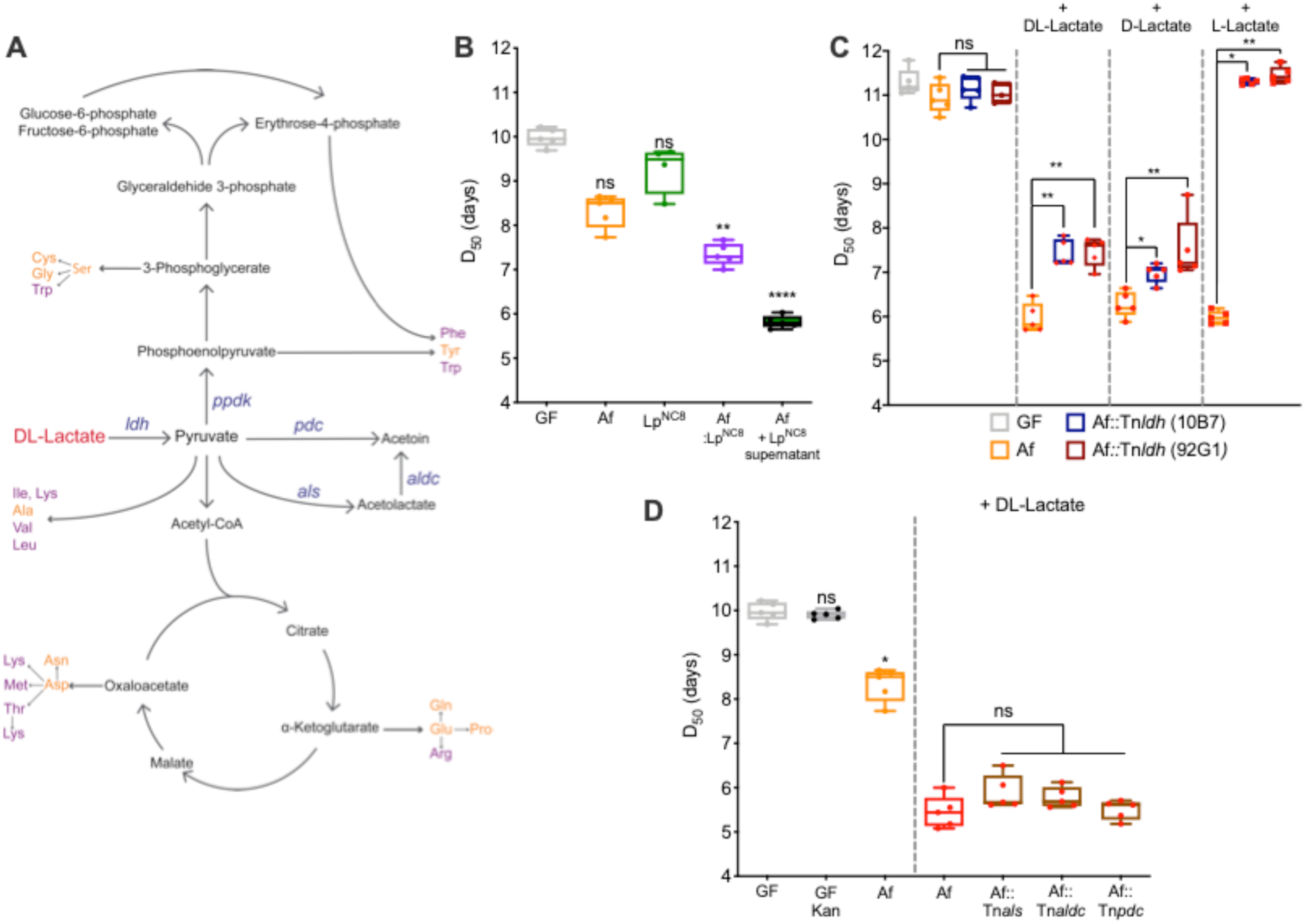
Lactate utilization by *Acetobacter* is central to its physiological response to Lp^NC8^ and enhanced benefit on host growth. (A) Schematic representation of the main metabolic routes of DL-lactate utilization by *Acetobacter* species. Purple: Fly’s essential amino acids, Yellow: Fly’s non-essential amino acids. Blue: genes related with lactate consumption. (B) Developmental timing of Germ Free (GF, grey) larvae or GF larvae inoculated with 10^5^ CFU of *A. fabarum*^DsW_054^ (Af, orange), Lp^NC8^ (green), both strains (Af:Lp^NC8^, purple) or Af supplemented with the supernatant from 72h culture of Lp^NC8^ (black, filled green) in complete HD. (C) Developmental timing of GF (grey) larvae or GF larvae inoculated with 10^5^ CFU of Af (orange), Af::Tn*ldh* (10B7) (blue) or Af::Tn*ldh* (92G1) (brown) supplemented with sterile PBS, DL-lactate, D-lactate or L-lactate in complete HD. (D) Developmental timing of GF (grey) larvae or GF larvae inoculated with 10^5^ CFU of Af (orange) or Af (red), Af:Tn*als* (brown), Af:Tn*aldc* (brown), Af:Tn*pdc* (brown) supplemented with DL-lactate in complete HD or complete HD supplemented with 50 µg/mL of Kanamycin (GF and Af mutants). Boxplots show minimum, maximum and median, each point represents a biological replicate. We performed Kruskal-Wallis test followed by uncorrected Dunn’s tests to compare each condition to the GF treated condition or the Af condition when indicated. ns: non-significant, *: p-value<0,05 **: p-value<0,005, ****: p-value<0,0001. See also Figure S5.

### *Acetobacter* acetoin pathway is not limiting for lactate mediated enhancement of *Acetobacter* larval growth promotion

After LDH conversion of lactate to pyruvate, acetoin can be produced from pyruvate either directly through the pyruvate decarboxylase (PDC) or by the successive action of the α-acetolactate synthase (ALS) and α-acetolactate decarboxylase (ALDC) with acetolactate as the intermediate product (Fig. 5A). To investigate if the acetoin production pathway is necessary to the lactate mediated enhancement of *Acetobacter* benefit to larvae development, we assessed the development of larvae mono-associated with each of the acetoin pathway mutants, Af::Tn*pdc*, Af::Tn*als*, Af::Tn*aldc* supplemented with DL-lactate. Of note, the mutants do not show any growth impairment on liquid HD (Fig. S5F) and previous analyses of these mutants showed that even if acetoin production is significantly reduced it is not fully inhibited; the Af::Tn*als* and Af::Tn*aldc* mutants produce 3 times less acetoin than Af and Af::Tn*pdc* in rich liquid media (Sommer and Newell, 2018). However, all the mutants in the genes responsible for acetoin production enhance larval development upon addition of DL-lactate to the same extent as the WT strain (Fig. 5D). Therefore, we conclude that acetoin production is not a limiting metabolic step in Af for the positive effect of lactate on the development of Af mono-associated larvae.

Another possible utilization of lactate by *Acetobacter* strains is the conversion from pyruvate to phosphoenolpyruvate (PEP) by the enzyme pyruvate phosphate dikinase (PPDK, Fig. 5A). PEP is a precursor for the synthesis of many cellular building blocks through the gluconeogenesis and the pentose phosphate pathways. We hypothesize that DL-lactate consumption by Af results in a higher flux towards biosynthetic pathways. However, Tn disruption of the *ppdk* gene has a strong effect on Af fitness in HD, completely precluding the growth of the mutant strains in this media (Fig. S5G) making it impossible to test them in our setting to obtain a complete genetic characterization of the phenotype.

### Lactate-dependent *Acetobacter* stimulation of larval growth evokes metabolites release enhancing host anabolic metabolism and resistance to oxidative stress

We next sought to characterize the molecular mechanisms involved in the enhancement of the growth promoting effect of *Acetobacter* strains upon lactate supplementation by a metabolic approach, using untargeted metabolomics (Fig. 6). To this end, we used Af as a model bacteria since it reproduces the phenotype of Ap^WJL^, and Af’s loss of function mutant Af::Tn*ldh* (clone 10B7). We capitalized on these two strains to characterize the bacterial metabolites produced at day 3 upon L-lactate supplementation in absence or presence of *Drosophila* larvae on HD (Fig. 6A, see Methods). We chose this time point to collect the samples because at day 3 post mono-association and lactate supplementation, we start observing significant larval size gains when compared to GF or *Acetobacter* mono-associated larvae. Also, at this time point larvae are actively increasing their size and mass and have not yet reached the critical weight to enter metamorphosis (Fig. 1B).

**Fig. 6:**
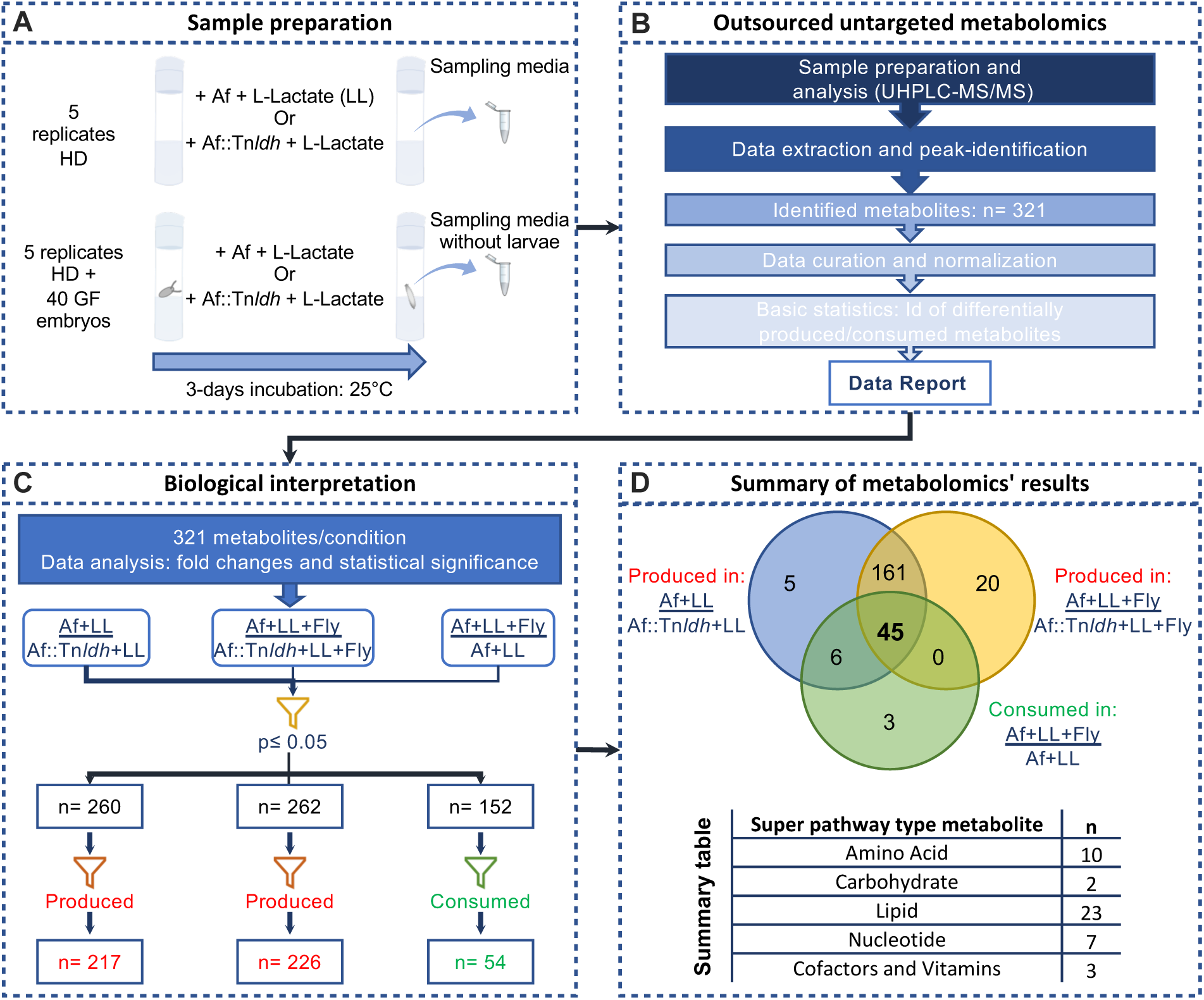
Lactate-dependent *Acetobacter* stimulation of larval growth evokes metabolites release enhancing host anabolic metabolism and resistance to oxidative stress. (A) Schematic representation of sample preparation for metabolomic analysis. (B) Outsourced untargeted metabolomics and data analysis pipeline. (C) Investigator driven data analysis and biological interpretation. (D) Venn diagram of the identified metabolites in the three test conditions. Our analysis points to 45 metabolites of interest belonging to all major metabolite families. See Table 1 for a detailed list of metabolites.

Untargeted metabolomic analyses based on Ultra High-Performance Liquid Chromatography coupled to Tandem Mass Spectrometry (UHPLC-MS/MS) identified 321 different metabolites (Fig. 6B). We first calculated the fold changes of the metabolites among four conditions: Af+LL/Af::Tn*ldh*+LL, Af+LL+Fly/Af::Tn*ldh*+LL+Fly and Af+LL+Fly/Af+LL (Fig. 6C, Table S2). As shown above, Af::Tn*ldh* fails to consume L-lactate and does not accelerate larval development (Fig. 5C and Fig S5D-E). Thus, the first two comparisons allow us to identify the differentially produced/consumed metabolites by Af upon L-lactate supplementation in absence or presence of the larvae, respectively. The third comparison, Af+LL+Fly/Af+LL, allow us to identify the metabolites that are produced/consumed by the larvae when they are mono-associated with Af and supplemented with L-lactate. From the three different sets of differentially produced/consumed metabolites, we selected only the metabolites that differed with statistical significance between experimental groups (Welch’s two-sample t-test, p≤0.05, Fig. 6C, Table S2). Next, we filtered the data sets in order to retain only the metabolites differentially produced by Af in absence or presence of the larvae upon L-lactate supplementation and the metabolites differentially consumed by the larvae in these conditions. The filtering generated three different sets of metabolites. The first is composed of 217 metabolites that are produced by Af upon L-lactate supplementation when growing on HD. The second comprises 226 metabolites that are produced by Af upon L-lactate supplementation when growing on HD in the presence of larvae. The third includes 54 metabolites that are consumed by larvae when mono-associated with Af and supplemented with L-lactate (Fig. 6C, Table S2). Finally, we crossed the three sets of metabolites in order to retain only the metabolites that are produced by Af upon L-lactate supplementation in both presence or absence of larvae and that at the same time are consumed by the larvae (Fig. 6D, venn diagram). These analyses provide us with a set of 45 metabolites encompassing all main metabolite families such as amino acids, carbohydrates, lipids, nucleotides, co-enzymes, cofactors and vitamins with a clear overrepresentation of amino acids derivatives and phospholipids (Fig. 6D, summary table and Table 1).

**Table 1:** Final metabolite candidate set, product of the analyses described in Fig. 6. Metabolites produced by Af upon L-lactate treatment in complete HD, in a *ldh* dependent manner and consumed by larvae.

| Super Pathway | Sub Pathway | Biochemical Name | Fold change (p<0.05) |  |  |
| --- | --- | --- | --- | --- | --- |
|  |  |  | <u>Af+LL</u><br>Af::Tn*ldh*+LL | <u>Af+LL+Fly</u><br>Af::Tn*ldh*+LL+Fly | <u>Af+LL+Fly</u><br>Af+LL |
| Amino Acid | Lysine Metabolism | pipecolate | 228.86 | 1119.99 | 0.79 |
|  | Tryptophan Metabolism | indoleacetate | 5.85 | 6.22 | 0.86 |
|  | Methionine, Cysteine, SAM and Taurine Metabolism | S-adenosylmethionine (SAM) | 2.94 | 1.64 | 0.56 |
|  |  | S-adenosylhomocysteine (SAH) | 3.26 | 2.11 | 0.65 |
|  |  | homocysteine | 9.84 | 7.28 | 0.65 |
|  |  | cysteine | 8.31 | 5.87 | 0.85 |
|  |  | S-methylcysteine | 14.10 | 7.95 | 0.54 |
|  | Polyamine Metabolism | spermidine | 373.79 | 92.21 | 0.25 |
|  | Glutathione Metabolism | cysteinylglycine | 4.22 | 1.43 | 0.34 |
|  |  | cys-gly, oxidized | 15.23 | 1.92 | 0.13 |
| Carbohydrate | Glycolysis, Gluconeogenesis, and Pyruvate Metabolism | dihydroxyacetone phosphate (DHAP) | 19.84 | 11.46 | 0.71 |
|  | Nucleotide Sugar | UDP-glucuronate | 3.86 | 1.94 | 0.50 |
| Lipid | Long Chain Monounsaturated Fatty Acid | eicosenoate (20:1) | 12.07 | 3.27 | 0.48 |
|  | Fatty Acid, Monohydroxy | 2-hydroxypalmitate | 9.68 | 4.99 | 0.42 |
|  |  | 2-hydroxystearate | 31.57 | 13.87 | 0.38 |
|  |  | 3-hydroxylaurate | 12.74 | 5.64 | 0.45 |
|  |  | 3-hydroxymyristate | 48.30 | 7.43 | 0.22 |
|  |  | 3-hydroxypalmitate | 105.93 | 34.11 | 0.25 |
|  |  | 3-hydroxystearate | 97.11 | 46.57 | 0.30 |
|  | Phosphatidylcholine (PC) | 1-palmitoyl-2-palmitoleoyl-GPC | 8.34 | 2.33 | 0.28 |
|  |  | 1-palmitoyl-2-oleoyl-GPC | 38.40 | 33.83 | 0.26 |
|  |  | 1-palmitoleoyl-2-oleoyl-GPC | 8.79 | 1.62 | 0.18 |
|  |  | 1-stearoyl-2-oleoyl-GPC | 67.53 | 17.79 | 0.26 |
|  |  | 1,2-dioleoyl-GPC | 135.60 | 276.71 | 0.32 |
|  | Phosphatidylethanolamine (PE) | 1,2-dipalmitoyl-GPE | 6.14 | 1.63 | 0.27 |
|  |  | 1-palmitoyl-2-oleoyl-GPE | 100.32 | 84.04 | 0.27 |
|  |  | 1-palmitoyl-2-linoleoyl-GPE | 10.83 | 1.91 | 0.18 |
|  |  | 1-stearoyl-2-oleoyl-GPE | 5.66 | 1.49 | 0.26 |
|  |  | 1,2-dioleoyl-GPE | 156.95 | 77.19 | 0.29 |
|  | Phosphatidylglycerol (PG) | 1,2-dipalmitoyl-GPG | 13.29 | 2.55 | 0.19 |
|  |  | 1-palmitoyl-2-oleoyl-GPG | 4.75 | 1.43 | 0.30 |
|  |  | 1-stearoyl-2-oleoyl-GPG | 4.10 | 1.38 | 0.34 |
|  |  | 1,2-dioleoyl-GPG | 3.55 | 1.35 | 0.38 |
|  | Sphingolipid Synthesis | sphinganine | 236.83 | 651.75 | 0.36 |
|  |  | hexadecasphinganine | 676.50 | 312.83 | 0.37 |
| Nucleotide | Purine Metabolism, Adenine containing | adenosine 5'-monophosphate (AMP) | 264.58 | 146.97 | 0.32 |
|  |  | N6-methyladenosine | 16.48 | 8.33 | 0.63 |
|  |  | guanosine 5'- monophosphate (5'-GMP) | 12.58 | 4.64 | 0.37 |
|  | Pyrimidine Metabolism, Orotate containing | dihydroorotate | 17.16 | 10.60 | 0.65 |
|  |  | uridine 5'-monophosphate (UMP) | 11.18 | 2.82 | 0.25 |
|  |  | 2'-deoxyuridine | 8.17 | 3.80 | 0.47 |
|  | Purine and Pyrimidine Metabolism | methylphosphate | 10.28 | 6.04 | 0.36 |
| Cofactors and Vitamins | Nicotinate and Nicotinamide Metabolism | nicotinamide adenine dinucleotide (NAD+) | 5.32 | 1.53 | 0.29 |
|  | Riboflavin Metabolism | flavin mononucleotide (FMN) | 3.24 | 1.42 | 0.44 |

The 45 differentially-produced metabolites constitute a large repertoire of molecules produced by *Acetobacter* upon lactate utilization and potentially accessible to the developing larvae. This particular combination of metabolites contains essential building-blocks and regulators for the host’s core anabolic process (nucleotides: AMP, GMP, UMP and cofactors/vitamins: NAD+, FMN, pyridoxamine phosphate) as well as regulator or intermediates of metabolic and developmental pathways (co-enzymes: SAM and SAH; phospholipids: biosynthetic intermediates of phosphatidylcholine, phosphatidylethanolamine and phosphatidylglycerol pathways and sphingolipids: sphinganine) and effectors of oxidative stress resistance (spermidine, cysteinylglycine). The collective action of these metabolites may converge to sustain linear larval growth and development despite a suboptimal nutritional environment. Altogether our work identifies a fruitful metabolic cooperation among commensal bacteria that support their physiology and would boost host juvenile growth while facing amino-acids scarcity.

## DISCUSSION

Here, we identify a beneficial metabolic dialogue among frequently co-habiting species of *Drosophila*’s commensal bacteria which optimizes host juvenile growth and enables cross-feeding and nutrient-provision upon chronic amino acids scarcity. Such benefit is also observed in full HDs containing optimal amino-acid contents as well as on fruit-based diets indicating that the metabolic cooperation among commensal bacteria and their influence on host growth is not restricted to artificial or poor nutritional conditions.

Using low amino acids containing HDs as an experimental model, we show that *L. plantarum* captures the essential amino-acids and B-vitamins synthetized by the *Acetobacter* species to fulfill its auxotrophic requirements. In parallel, *Acetobacter* species exploit the lactate produced by *L. plantarum* as an additional carbon source that alters its metabolic state and physiology. Such metabolic interactions support an optimized growth of both commensal species in the diet and an increased colonization of the host.

Previous work has shown a positive correlation between host-associated microbial counts and linear larval growth on *Drosophila* (Keebaugh et al., 2019). Moreover, inert microbial biomass (heat-killed microbes) can accelerate larval development (Bing et al., 2018; Storelli et al., 2011) and impact *Drosophila* lifespan (Yamada et al., 2015). Here, we show that the metabolic cooperation between Ap^WJL^ and Lp^NC8^ increases bacterial biomass in the nutritional substrate, which slightly increase larval growth. However, the bacterial biomass alone never reproduces to the same extent the positive impact of live Ap^WJL^:Lp^NC8^ bi-association or lactate supplemented Ap^WJL^ mono-association on host growth. Instead, we show that lactate utilization by *Acetobacter* species rewires its carbon metabolism resulting in the enhanced and *de novo* production of a panoply of anabolic metabolites that would support enhanced host systemic growth.

Studies have previously shown that cooperation among the gut microbes can influence other aspects of *Drosophila* physiology. For example, multiple fermentation products of *L. brevis* foster the growth of *A. fabarum* on a fly diet leading to depletion of dietary glucose, consequently triggering reduced TAG levels in the adult host (Newell and Douglas, 2014; Sommer and Newell, 2018). Moreover, multi-microbe interactions among the *Acetobacter*, *Lactobacillus* species and yeast were shown to influence additional adult traits such as olfaction and egg laying behavior (Fischer et al., 2017), food choice behavior (Leitão-Gonçalves et al., 2017), lifespan and fecundity (Gould et al., 2018) and immunity (Fast et al., 2020). Therefore, along with these studies, our work provides an entry point to further deepen the understanding of how metabolites originating from microbial metabolic networks shape the biology of their host.

In this study, we confirm that lactate is a key metabolite supporting the metabolic crosstalk between different microbial species. Lactate supplementation to *Acetobacter* species triggers the release of metabolic by-products that include ribonucleotides AMP, GMP and UMP, vitamin and amino-acid derivatives: SAM, SAH, NAD+, FMN and pyridoxamine phosphate, which are co-factors for enzymes involved in multiple host metabolic pathways. These metabolites are essential for optimal larval growth and survival (Consuegra et al., 2020; Mishra et al., 2018; Sang, 1956). Fatty acids and membrane lipids are another group of metabolites whose production is enhanced by lactate presence. Among this group, we found mostly phospholipids such as phosphatidylcholine (PC), phosphatidylethanolamine (PE) and phosphatidylglycerol (PG) and a sphingolipid precursor, sphinganine. In *Drosophila*, PE, PC, PG and sphingolipids are part of the membrane phospholipids repertoire, with PE being the largely dominating species (Carvalho et al., 2012). Previously it was established that the total content of membrane lipids increases during larval growth, until a clear pause that occurs in the third instar just prior to the time when larvae stop feeding and enter the wandering stage. This indicates that feeding larvae favor new membrane synthesis and tissue growth over lipid storage (Carvalho et al., 2012). In the same study, it was shown that dietary lipids directly influence membrane lipids proportions, including phospholipids and sphingolipids. In mammals, sphingolipid balance has a central role in controlling nutrient utilization and growth (Holland et al., 2007). Sphingolipids are also activators of serum response element binding protein signaling, which controls biosynthesis of fats (Worgall, 2008). Despite a relatively smaller literature on *Drosophila* sphingolipids, these lipids seem as critical to developmental and metabolic processes in the fly as they are to mammals (Kraut, 2011). Although *Drosophila* cells can synthesize *de novo* all the fatty acids for survival, they incorporate different dietary lipids into the membrane lipids if found in the diet (Carvalho et al., 2012). Therefore, we propose that larvae preferentially utilize the PC, PE, PG and sphingolipids intermediates produced by *Acetobacter* species upon lactate utilization to foster membrane synthesis, tissue growth, and metabolic processes such as lipid storage and response to nutrient availability.

Lactate utilization also triggers another major class of metabolites released by *Acetobacter* species that confers oxidative stress resistance. Specifically, we found cysteinylglycine and spermidine. Cysteinylglycine is an intermediate of glutathione (GSH) metabolism, the most abundant cellular antioxidant (Forman et al., 2009). It is produced by GSH hydrolysis or by action of the enzyme γ-L-Glutamyl-transpeptidase (GGT). GGT transfers the γ-glutamyl group of GSH onto amino acids forming γ-glutamyl peptides and cysteinylglycine. These intermediaries can be recycled and used to resynthesize GSH and maintain its cellular pool, which protects cells from oxidative damage and maintains redox homeostasis (Ursini et al., 2016). Of note, during *Drosophila* larval development, in addition to its antioxidant role, GSH also contributes to ecdysteroid biosynthesis including the biologically active hormone 20-hydroxyecdysone which plays an essential role in promoting juvenile growth and maturation (Enya et al., 2017). Spermidine is a natural polyamine widely found in both prokaryotes and eukaryotes including flies and mammals. Nutritional supplementation of spermidine increases the lifespan of yeast, worms, flies and human cells through inhibition of oxidative stress (Eisenberg et al., 2009). The mode of action of spermidine, mainly through autophagy regulation, is emerging but evidence for other mechanisms exist such as inflammation reduction, lipid metabolism, and regulation of cell growth, proliferation and death (Minois, 2014; Minois et al., 2012). Oxidative stress resistance in *Drosophila* has been largely reported to improve adult physiology including lifespan extension. We therefore posit that larvae’s physiology and growth potential are also supported by such metabolites obtained from their microbial partners, especially during development on a suboptimal diet. Further work, including testing individual metabolites and their combinations, will be required to identify the specific compounds or cocktails produced by *Acetobacter* upon lactate utilization supporting acceleration of larval development.

Beyond essential nutrient provision and metabolic cooperation between commensals and their host, we posit that other bacteria mediated mechanisms would also contribute to enhanced host growth. Indeed, upon lactate utilization *Acetobacter* may release molecules which would activate host endocrine signals and promote anabolism. Accordingly, it was recently shown that acetate produced by *Acetobacter* improves larval growth by impacting host lipid metabolism through the activation of the IMD signalling pathway in entero-endocrine cells and the release of the endocrine peptide tachykinin (Kamareddine et al., 2018). However, this mechanism is unlikely to be at play here due to the high content of acetate in our fly diet.

Collectively our results deconstruct the intertwined metabolic networks forged between commensal bacteria that support juvenile growth of the host. This work contributes to the understanding of how the microbiota activities as a whole, influence host nutritional and metabolic processes supporting host juvenile growth despite a stressful nutritional environment.

## LIMITATIONS OF THE STUDY

The complete genetic characterization of Lactate-dependent *Acetobacter* stimulation of larval growth was hampered by the lethality of *Acetobacter* mutants affecting the central metabolic pathways while growing in complete HD. Instead, using metabolomics, we pinpoint a large repertoire of molecules produced by *Acetobacter* upon lactate utilization and accessible to the developing larvae. Further studies will be necessary to test the 45 candidate metabolites, individually or in combinations, to identify the minimal metabolite cocktail enhancing the development of GF larvae or larvae mono-associated with *Acetobacter*. Moreover, functional analyses in the host would be required to identify the metabolic pathways sustained by commensal bacteria and involved in the anabolic growth of the host.

## METHODS

All methods can be found in the accompanying Transparent Methods supplemental file.

## RESOURCE AVAILABILITY

### Lead Contact

Further information and requests for resources should be addressed to the Lead Contact, François Leulier.

### Materials availability

This study did not generate new unique reagents.

### Data and code availability

Table 1 and Table S2 provide the main results derived from the metabolomic analysis presented in this study.

## ACKNOWLEDGEMENTS

We would like to thank Dali Ma for critical reading and editing of the manuscript and valuable suggestions, John Chaston and Peter Newell for *Acetobacter fabarum* strains and mutants, the ArthroTools platform of the SFR Biosciences (UMS3444/US8) for fly equipment and facility. Research in F.L’s lab is supported by the ENS de Lyon, CNRS and the FINOVI foundation. Research in P.d.S lab is supported by INRA and INSA Lyon. J.C. is funded by a postdoctoral fellowship from the “Fondation pour la Recherche Médicale” (FRM, SPF20170938612). T.G. is funded by a Ph.D fellowship from ENS de Lyon.

## AUTHOR CONTRIBUTIONS

Conceptualization, J.C. and F.L.; Methodology, J.C. and F.L.; Validation, J.C. and F.L.; Formal Analysis, J.C.; Investigation, J.C., T.G., H.A., H.G., I.R. and P. daS.; Data Curation, J.C.; Writing – Original Draft, J.C. and F.L.; Writing – Review & Editing, J.C. T.G., P.daS., and F.L.; Visualization, J.C.; Supervision, J.C. and F.L.; Project Administration, J.C. and F.L.; Funding Acquisition, F.L.

