## Supplemental Information for "Metabolic cooperation among commensal bacteria supports *Drosophila* juvenile growth under nutritional stress"

#### SUPPLEMENTAL FIGURES

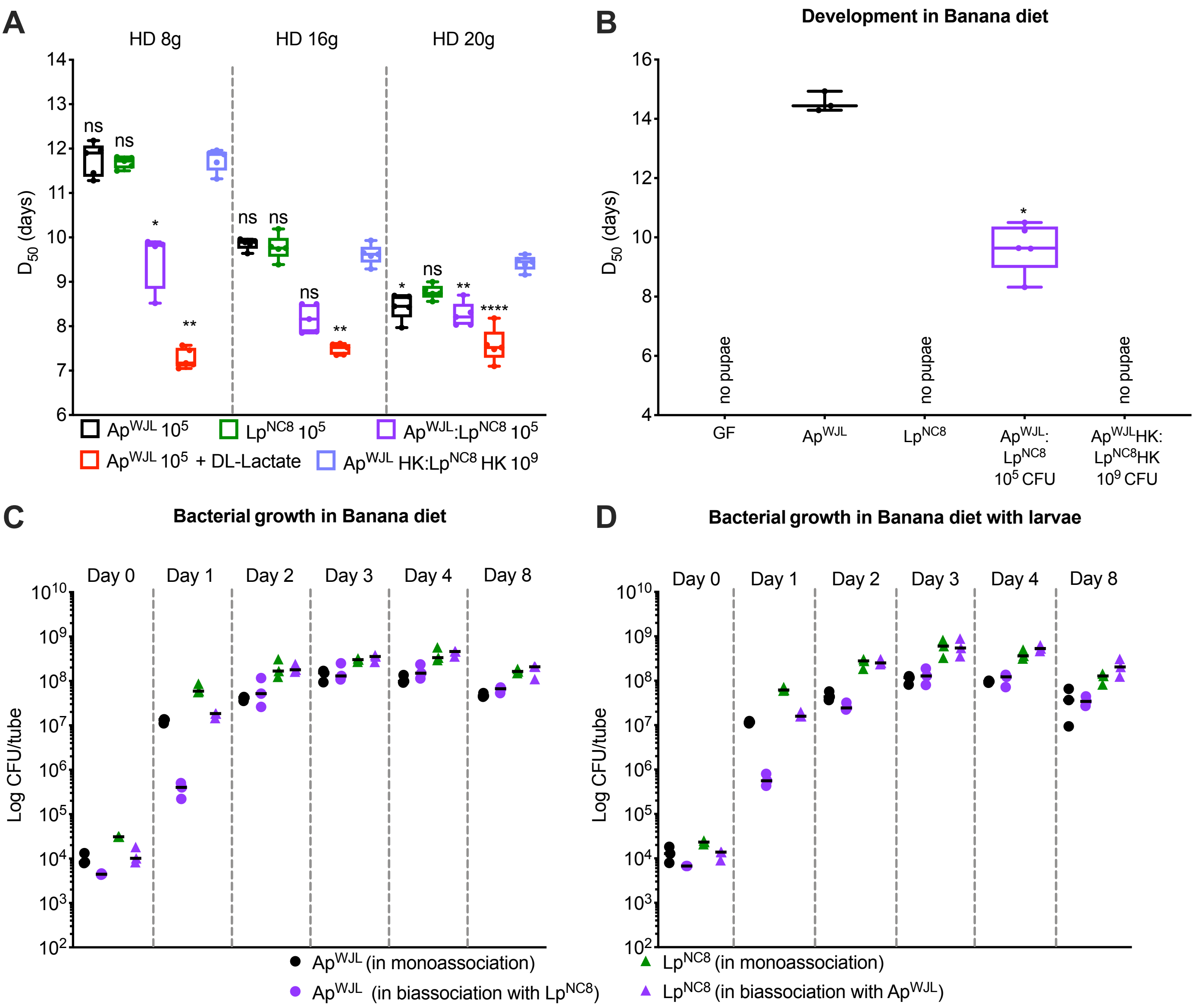


**Fig. S1 (related to Fig. 1 and Fig. 3):** (A-B) Developmental timing of Germ Free (GF, grey) larvae or GF larvae inoculated with $10^5$ CFU of live $Ap^{WJL}$ (black), $Lp^{NC8}$ (green), $Ap^{WJL}$:$Lp^{NC8}$ bi-association (purple) or $10^9$ CFU heat-killed $Ap^{WJL}$:$Lp^{NC8}$ bi-association (light purple) in HD with a total amino acid content of 8 g/L, 16 g/L, or 20 g/L (A) or Banana-diet (B). (C-D) Load of $Ap^{WJL}$ and $Lp^{NC8}$ in mono- (black and green, respectively), bi-association (purple) or $Ap^{WJL}$ mono-association supplemented with DL-lactate at a final concentration of 0.6 g/L (red) in solid Banana-diet with (D) and without (C) larvae, from day 0 to 4 days and 8 days after inoculation. Boxplots show minimum, maximum and median. Points represent biological replicates. We performed Kruskal-Wallis test followed by uncorrected Dunn's tests to compare each condition to the GF treated condition. ns: non-significant, *: p-value<0,05, **: p-value<0,005, ***: p-value<0,0005 ****: p-value<0,0001. Dot plots shows mean and each dot represents an independent biological replicate.

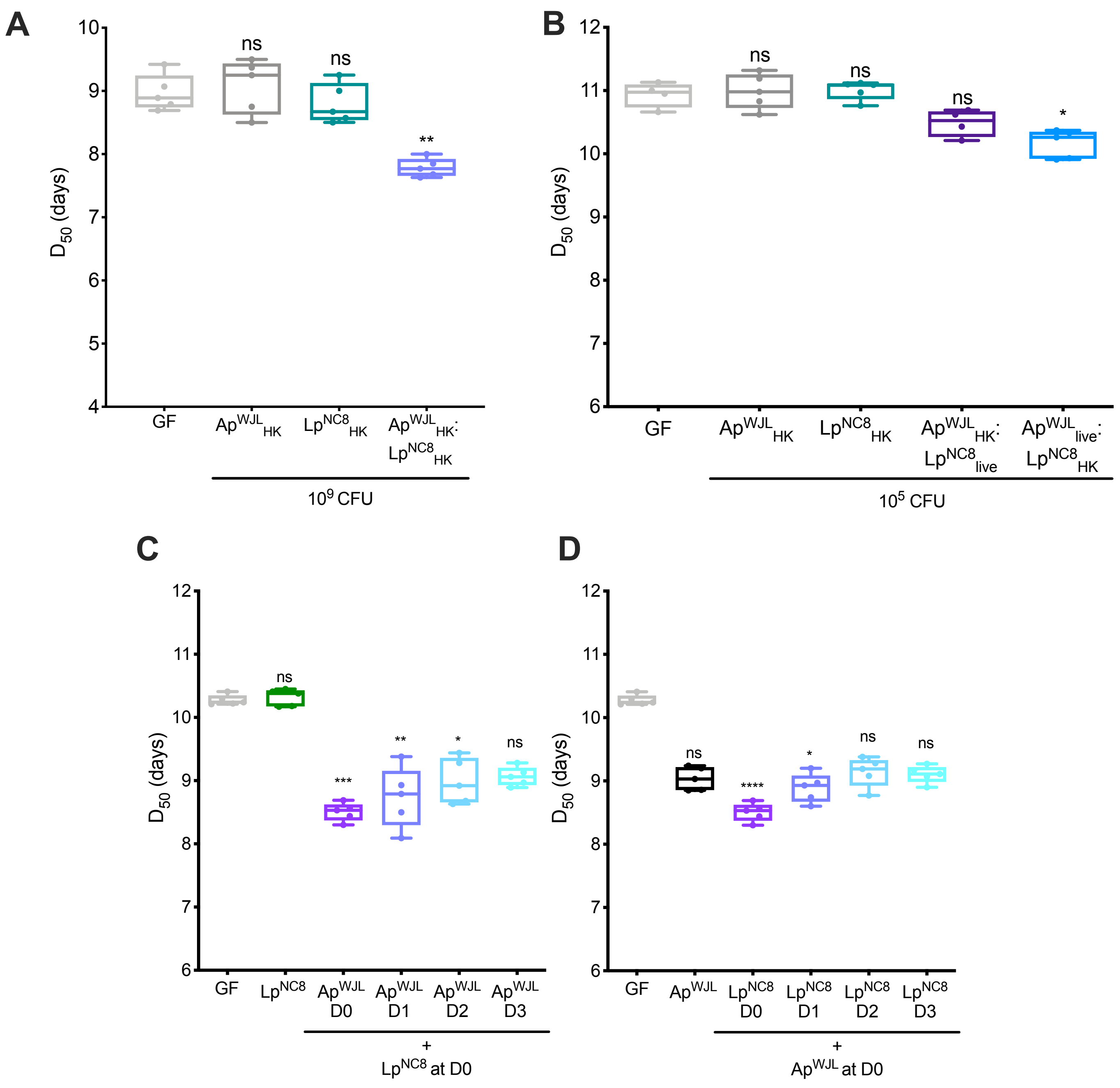


**Fig. S2 (related to Fig. 1):** (A) Developmental timing of Germ Free (GF, light grey) larvae or GF larvae inoculated with $10^9$ CFU of $Ap^{WJL}$ Heat-Killed ($Ap^{WJL}_{HK}$, dark gray), $Lp^{NC8}_{HK}$ (turquoise) or $Ap^{WJL}_{HK}$: $Lp^{NC8}_{HK}$ bi-association (light purple). (B) Developmental timing of GF (light grey) larvae or GF larvae inoculated with $10^5$ CFU of $Ap^{WJL}_{HK}$ (dark gray), $Lp^{NC8}_{HK}$ (turquoise), $Ap^{WJL}_{HK}$ plus live $Lp^{NC8}$ ($Ap^{WJL}_{HK}$:$Lp^{NC8}_{live}$, dark purple) or $Ap^{WJL}_{live}$:$Lp^{NC8}_{HK}$, light blue). (C) Developmental timing of GF (grey) larvae or GF larvae inoculated with $10^5$ CFU of $Lp^{NC8}$ at D0 and subsequently at D0/1/2/3 with ~$10^5$ CFU of $Ap^{WJL}$. (D) Developmental timing of GF (grey) larvae or GF larvae inoculated with $10^5$ CFU of $Ap^{WJL}$ at D0 and subsequently at D0/1/2/3 with $10^5$ CFU of $Lp^{NC8}$. Boxplots show minimum, maximum and median. Points represent biological replicates. We performed Kruskal-Wallis test followed by uncorrected Dunn's tests to compare each condition to the GF treated condition. ns: non-significant, *: p-value<0,05, **: p-value<0,005, ***: p-value<0,0005 ****: p-value<0,0001.

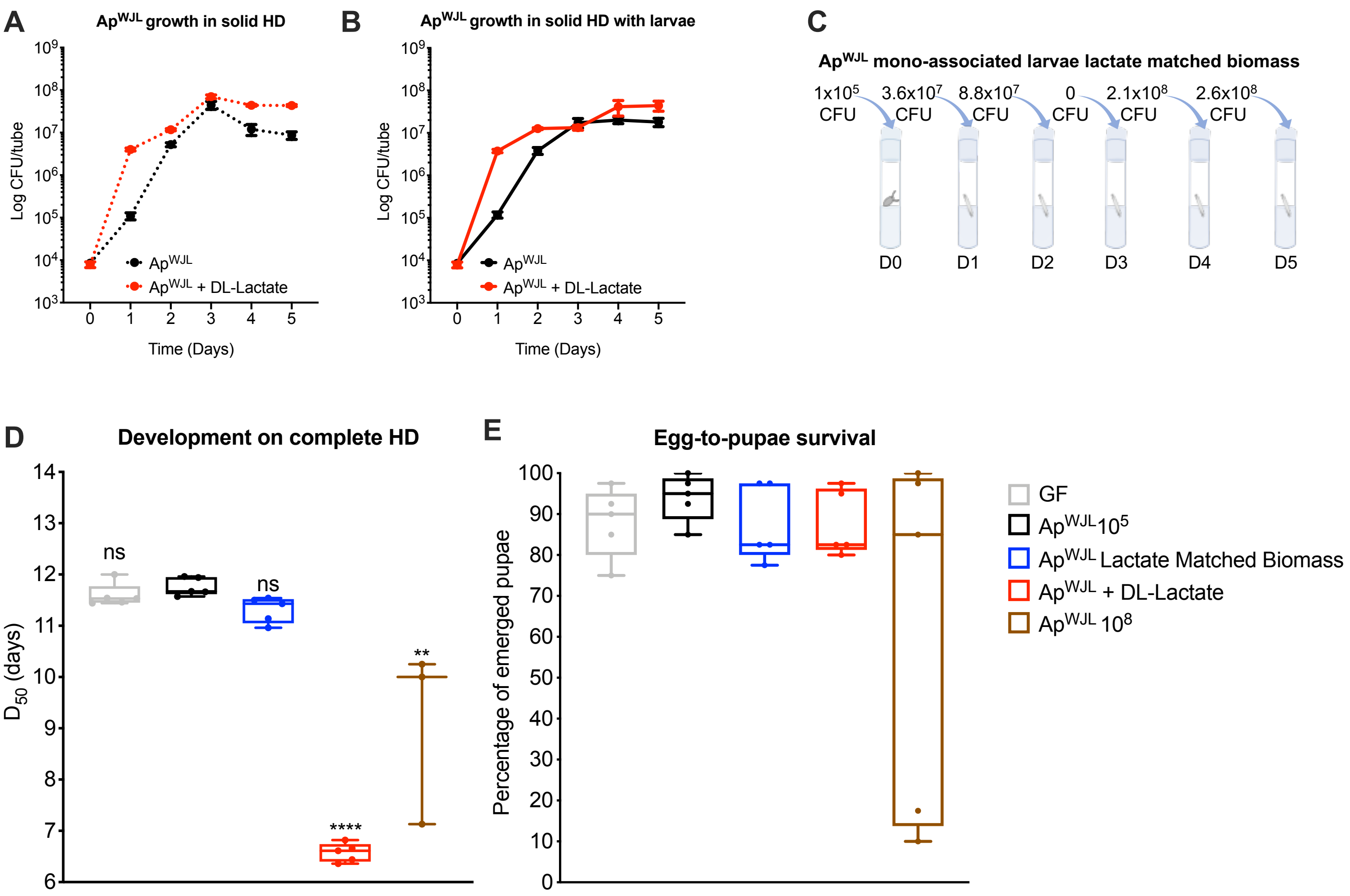


**Fig. S3 (related to Fig. 3):** (A-B) Load of Ap$^{WJL}$ in solid HD supplemented (red) or not (black) with DL-lactate at a final concentration of 0.6 g/L with (B) and without (A) larvae, from day 0 to 5 days after inoculation. (C) Graphical representation of the daily Ap$^{WJL}$ biomass supplementation to Ap$^{WJL}$ mono-associated larvae in order to match the biomass reached upon DL-lactate supplementation, according with Fig. S3B. (D) Developmental timing of Germ Free (GF, light grey) larvae or GF larvae inoculated with $10^5$ CFU of Ap$^{WJL}$ (black) or Ap$^{WJL}$ mono-associated larvae supplemented daily with live Ap$^{WJL}$ biomass (blue) or DL-lactate (red) and GF larvae inoculated with $10^8$ CFU of Ap$^{WJL}$ (brown). (E) Percentage of the emerged pupae from the developmental timing experiment of Fig. S3D. Symbols represent the means ± SEM of three biological replicates except for panel (A-B). Boxplots show minimum, maximum and median. Points represent biological replicates. We performed Kruskal-Wallis test followed by uncorrected Dunn's tests to compare each condition to the GF treated condition. ns: non-significant, *: p-value<0,05, **: p-value<0,005, ***: p-value<0,0005 ****: p-value<0,0001.

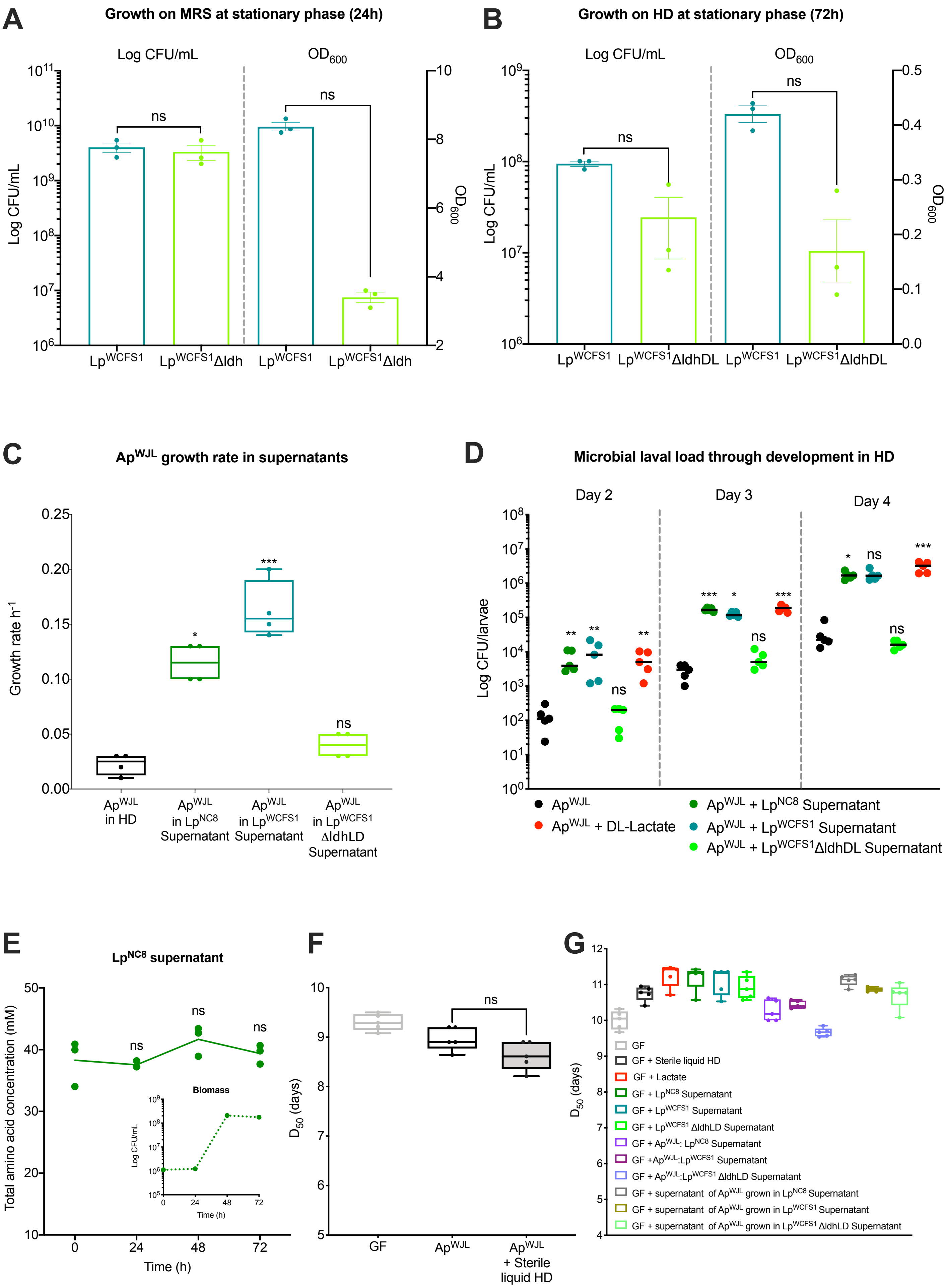
A
Growth on MRS at stationary phase (24h)
Log CFU/mL
OD600
ns
ns
Log CFU/mL
OD600
LpWCFS1
LpWCFS1Δldh
LpWCFS1
LpWCFS1Δldh
B
Growth on HD at stationary phase (72h)
Log CFU/mL
OD600
ns
ns
Log CFU/mL
OD600
LpWCFS1
LpWCFS1ΔldhDL
LpWCFS1
LpWCFS1ΔldhDL
C
ApWJL growth rate in supernatants
Growth rate h-1
*
***
ns
ApWJL in HD
ApWJL in LpNC8 Supernatant
ApWJL in LpWCFS1 Supernatant
ApWJL in LpWCFS1 ΔldhDL Supernatant
D
Microbial laval load through development in HD
Day 2
Day 3
Day 4
Log CFU/larvae
ApWJL
ApWJL + DL-Lactate
ApWJL + LpNC8 Supernatant
ApWJL + LpWCFS1 Supernatant
ApWJL + LpWCFS1ΔldhDL Supernatant
E
LpNC8 supernatant
Total amino acid concentration (mM)
Time (h)
ns
ns
ns
Biomass
Log CFU/mL
Time (h)
F
D50 (days)
ns
GF
ApWJL
ApWJL + Sterile liquid HD
G
D50 (days)
GF
GF + Sterile liquid HD
GF + Lactate
GF + LpNC8 Supernatant
GF + LpWCFS1 Supernatant
GF + LpWCFS1 ΔldhLD Supernatant
GF + ApWJL: LpNC8 Supernatant
GF +ApWJL:LpWCFS1 Supernatant
GF + ApWJL:LpWCFS1 ΔldhLD Supernatant
GF + supernatant of ApWJL grown in LpNC8 Supernatant
GF + supernatant of ApWJL grown in LpWCFS1 Supernatant
GF + supernatant of ApWJL grown in LpWCFS1 ΔldhLD Supernatant

**Fig. S4 (related to Fig. 3):** (A-B) CFU count and $OD_{600}$ of $Lp^{WCSF1}$ (turquoise) or $Lp^{WCFS1}\Delta$ldhDL (light green) cultures at stationary phase in MRS (24h, A) or complete holidic diet (72h, B). Bars represent mean ± SEM. We performed Mann-Whitney test to compare OD and CFU counts of $Lp^{WCFS1}$ to $Lp^{WCFS1}\Delta$ldhDL. (C) Growth rate of $Ap^{WJL}$ on complete HD (black), $Lp^{NC8}$ supernatant (green), $Lp^{WCFS1}$ supernatant (turquoise) or $Lp^{WCFS1}\Delta$*ldhDL* supernatant (light green). We performed Mann-Whitney test to compare the growth rate of $Ap^{WJL}$ monoculture in HD to to the growth rate of $Ap^{WJL}$ growing in the supernatant of interest. (D) $Ap^{WJL}$ larval loads on complete HD (black) or complete HD supplemented with $Lp^{NC8}$ supernatant (green), $Lp^{WCFS1}$ supernatant (turquoise), $Lp^{WCFS1}\Delta$*ldhDL* supernatant (light green) or DL-lactate at a final concentration of 0.6 g/L (red). We performed Kruskal-Wallis test followed by uncorrected Dunn's tests to compare each condition to the $Ap^{WJL}$ condition (E) HPLC quantification of total amino acid concentration (µM) in $Lp^{NC8}$ supernatant during growth in liquid HD. Inner panel: $Lp^{NC8}$ growth. Dot plots show mean and each point represent a biological replicate. We performed Kruskal-Wallis test followed by uncorrected Dunn's tests to compare each time point to T0. (F) Developmental timing of Germ Free (GF, light grey) larvae or GF larvae inoculated with $10^5$ CFU of $Ap^{WJL}$ supplemented (black, grey filling) or not (black) with 300 µL of sterile liquid HD. We performed Mann-Whitney test to compare the $D_{50}$ of $Ap^{WJL}$ to $Ap^{WJL}$ supplemented with steril HD. (G) Developmental timing of GF (grey) larvae or GF larvae supplemented with pure lactate (red), 300 µL of sterile liquid HD (black) or 300 µL of the different culture supernatants. ns: non-significant, *: p-value<0,05, **: p-value<0,005, ***: p-value<0,0005.

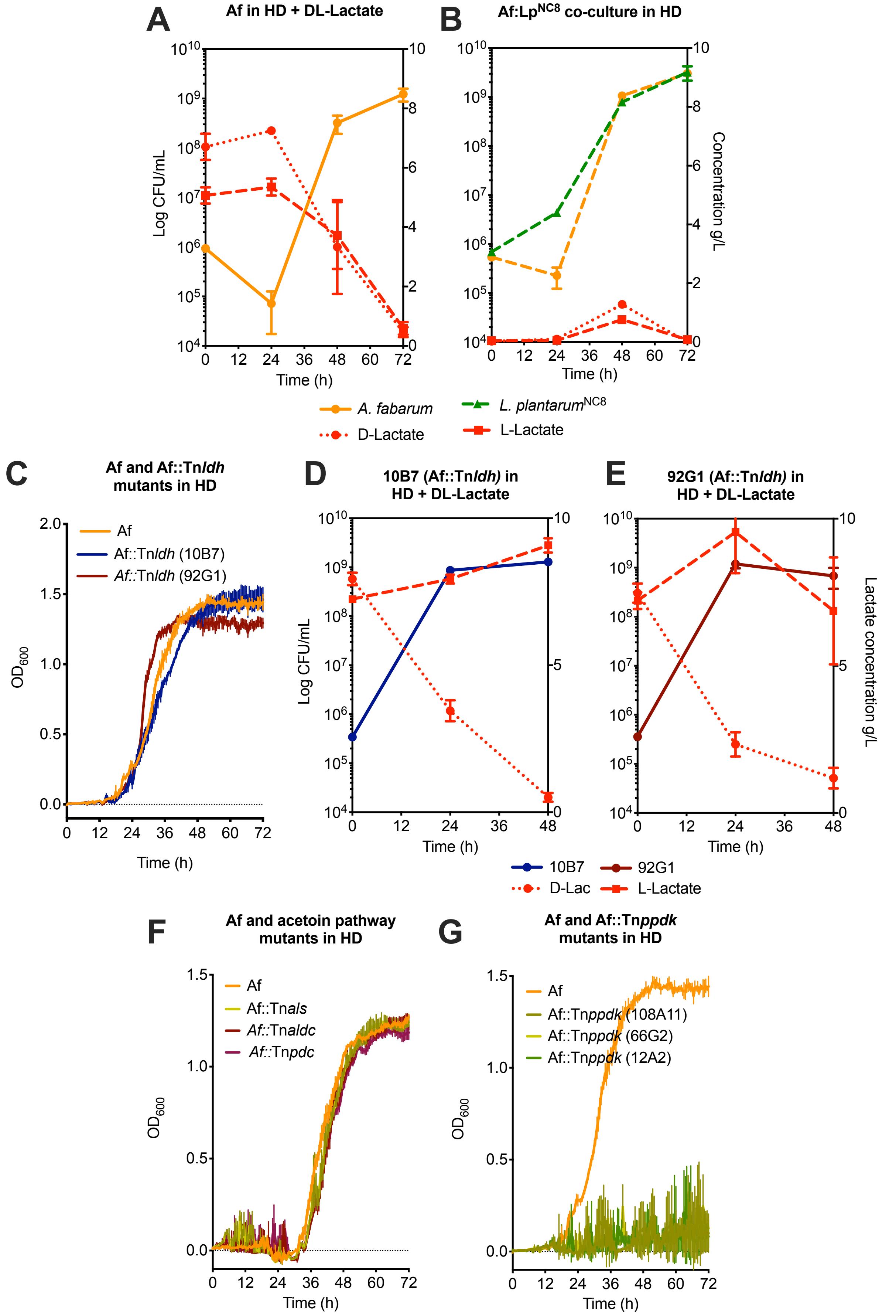


**Fig. S5 (related to Fig. 5):** (A) Growth curve of *A. fabarum* (orange) in liquid HD supplemented with DL-lactate. D- (dotted line) and L-lactate (dashed line) consumption (red) was quantified. (B) Growth curves in liquid HD of $Lp^{NC8}$ (dashed green line) or Af (dashed orange line) in co-culture with the respective D- (dotted red line) or L-lactate (dashed red line) levels (red). (C) Growth curve of *A. fabarum* (orange), Af::Tn*ldh* (10B7) (blue) or Af::Tn*ldh* (92G1) (brown) in liquid HD. (D-E) Growth curves in liquid HD supplemented with DL-lactate of Af::Tn*ldh* (10B7) (D) (blue line) or Af::Tn*ldh* (92G1) (E) (brown line) with the respective D- (dotted red line) or L-lactate (dashed red line) levels (red). (F) Growth curves of *A. fabarum* (orange), Af::Tn*als* (light green), Af::Tn*aldc* (brown) or Af::Tn*pdc* (dark red) in liquid HD. (G) Growth curves of *A. fabarum* (orange), Af::Tn*ppdk* (108A11) (green), Af::Tn*ppdk* (66G2) (light green) or Af:Tn*ppdk* (12A2) (dark green) in liquid HD. Symbols represent the means ± SEM of three biological replicates.

#### SUPPLEMENTAL TABLES

**Table S1:** Final amino acid concentration supplemented to complete HD in the amino acid cocktail supplementation experiment (Fig. 4)

| Amino acid | AA Mix (mg/L) | | |
|---|---|---|---|
| | **Ap @48h** | **Ap + Lactate @24h** | **Ap + Lactate @48h** |
| Arg | - | 36.802 | 1.337 |
| His | 7.467 | 39.088 | 12.911 |
| Ile | - | 14.913 | - |
| Leu | 9.091 | 19.230 | - |
| Lys | - | 71.720 | 33.243 |
| Met | - | - | - |
| Phe | 16.911 | 56.140 | 25.482 |
| Thr | 15.223 | 41.586 | 26.157 |
| Val | 5.115 | 22.169 | 17.314 |
| Ala | - | 21.621 | 80.409 |
| Asp | 0.241 | 13.920 | - |
| Glu | 4.703 | - | - |
| Gly | - | 8.284 | - |
| Pro | - | - | - |
| Ser | 18.407 | 38.394 | - |
| Tyr | - | 77.555 | 26.883 |
| **Total** | **77.160** | **461.428** | **223.740** |

**Table S3**. Strains used in this study

| Strain | Abbreviation | Genotype | Reference |
|---|---|---|---|
| *Acetobacter pomorum*$^{WJL}$ | Ap$^{WJL}$ | WT | Shin et al. 2011 |
| *Lactobacillus plantarum*$^{NC8}$ | Lp$^{NC8}$ | WT | Axelsson L et al. 2012 |
| *L. plantarum*$^{WCFS1}$ | Lp$^{WCFS1}$ | WT | Ferain et al. 1996 |
| *L. plantarum*$^{WCFS1}$ Δ*ldhDL* | Lp$^{WCFS1}$ΔldhDL | Δ*ldhDL* | |
| *A. fabarum*$^{DsW\_054}$ | Af | WT | Winans et al. 2017 |
| *A. fabarum*$^{DsW\_054}$ Tn::*ldh* (10B7) | Af::Tn*ldh* (10B7) | Tn::*ldh* | Winans et al. 2017 and Sommer & Newell 2018 |
| *A. fabarum*$^{DsW\_054}$ Tn::*ldh* (92G1) | Af::Tn*ldh* (92G1) | Tn::*ldh* | |
| *A. fabarum*$^{DsW\_054}$ Tn::*als* | Af::Tn*als* | Tn::*als* | |
| *A. fabarum*$^{DsW\_054}$ Tn::*aldc* | Af::Tn*aldc* | Tn::*aldc* | |
| *A. fabarum*$^{DsW\_054}$ Tn::*pdc* | Af::Tn*pdc* | Tn::*pdc* | |
| *A. fabarum*$^{DsW\_054}$ Tn::*ppdk* (108A11) | Af::Tn*ppdk* (108A11) | Tn::*ppdk* | |
| *A. fabarum*$^{DsW\_054}$ Tn::*ppdk* (66G2) | Af::Tn*ppdk* (66G2) | Tn::*ppdk* | |
| *A. fabarum*$^{DsW\_054}$ Tn::*ppdk* (12A2) | Af::Tn*ppdk* (12A2) | Tn::*ppdk* | |

### TRANSPARENT METHODS

#### *Drosophila* diets, stocks and breeding

*Drosophila* stocks were reared as described previously (Erkosar et al., 2015). Briefly, flies were kept at 25°C with 12/12-hour dark/light cycles on a yeast/cornmeal medium containing 50 g/L of inactivated yeast, 80 g/L of cornmeal, 7.4 g/L of agar, 4 mL/L of propionic acid and 5.2 g/L of nipagin. Germ-free stocks were established as described previously (Erkosar et al., 2014) and maintained in yeast/cornmeal medium supplemented with an antibiotic cocktail composed of kanamycin (50 µg/mL), ampicillin (50 µg/mL), tetracycline (10 µg/mL) and erythromycin (5 µg/mL). Axenicity was tested by plating fly media on nutrient agar plates. *Drosophila yw* flies were used as the reference strain in this work.

Experiments were performed on Holidic Diet (HD) without preservatives. Complete HD, with a total of 8 g/L, 16 g/L or 20 g/L of amino acids, were prepared as described by Piper at al. using the fly's exome matched amino acid ratios (FLYAA) (Piper et al., 2017). Briefly, sucrose, agar, amino acids with low solubility (Ile, Leu and Tyr) as well as stock solutions of metal ions and cholesterol were combined in an autoclavable bottle with milli-Q water up to the desired volume, minus the volume of solutions to be added after autoclaving. After autoclaving at 120°C for 15 min, the solution was allowed to cool down at room temperature to ~60 °C. Acetic acid buffer and stock solutions for the essential and non-essential amino acids, vitamins, nucleic acids and lipids precursors were added. Single nutrient deficient HD (Fig. 2 and Fig. 3C) were prepared following the same recipe excluding the nutrient of interest (named HDΔX, X being the nutrient omitted) as described in (Consuegra et al., 2020). Tubes used to pour the HD were sterilized under UV for 20 min. HD was stored at 4°C until use, for no longer than one week.

Banana diet was prepared with 200 mL of mixed banana, 300 mL of water and 3.5 g of agar. After autoclaving at 120°C for 15 min, 10 mL of diet were poured into UV-sterilized tubes. Banana diet was stored at 4°C and used the next day.

**Bacterial strains and growth conditions**

Strains used in this study are listed in Table S3. *A. pomorum* was cultured in 10 mL of Mannitol Broth (Bacto peptone 3 g/L, yeast extract 5 g/L, D-mannitol 25 g/L) in 50 mL flask at 30°C under 180 rpm agitation during 24h. *A. fabarum* strains were cultured in 10 mL of YPD (yeast extract 10 g/L, Bacto peptone 10 g/L, Glucose 8 g/L) in 50 mL flask at 30°C under 180 rpm agitation during 24h. *L. plantarum* strains were cultured in 10 mL of MRS broth (Carl Roth, Germany) in 15 mL culture tubes at 37°C, without agitation, overnight. Liquid or solid cultures of Af::Tn were supplemented with kanamycin (Sigma-Aldrich, Germany) at a final concentration of 50 µg/mL. CFU counts were performed for all strains on MRS agar (Carl Roth, Germany). For selective isolation of *Acetobacter* or *Lactobacillus* during cocultures or bi-association, MRS plates were supplemented with ampiciline (10 µg/mL) or kanamycin (50 µg/mL), respectively. Appropriated dilutions were plated using the Easyspiral automatic plater (Intersciences, Saint Nom, France). The MRS agar plates were then incubated for 24-48h at 30°C for *Acetobacter* strains or 37°C for Lactobacillus. CFU counts were done using the automatic colony counter Scan1200 (Intersciences, Saint Nom, France) and its counting software.

**Bacterial growth in liquid HD**

To assess bacterial growth in the fly nutritional environment we used a recently developed liquid HD comprising all HD components except agar and cholesterol (Consuegra et al., 2020). Liquid HD was prepared as described for solid HD. Single nutrient deficient liquid HD was prepared following the same recipe excluding the nutrient of interest. After growth in rich media, the strain to be tested was washed with PBS twice and inoculated at a final concentration of ~$10^6$ CFU/mL. For cocultures, the strains were inoculated in a 1:1 ratio. For growth assessment in microplates, 200 µL of media were inoculated in triplicate. Cultures were incubated in 96-well microtiter plates (Nunc™ Edge 2.0. Thermo Fisher Scientific) at 30°C for 72h. Growth was monitored using an SPECTROstar$^{Nano}$ (BMG Labtech GmbH, Germany) by measuring the optical density at 600 nm ($OD_{600}$) every 30 minutes. For growth assessment in flasks, 10mL of complete or single nutrient deficient HD were inoculated in triplicate. Cultures were incubated in 50 mL flasks at 30°C under 180 rpm during 72h. Bacterial growth was assessed by plating appropriated dilutions of the cultures every 24h on MRS agar as described above. In figures representing growth in flasks the symbols represent the means with standard error based on three biological replicates. Growth rates were computed by calculating the slope of the curve during exponential growth using SPECTROstar$^{Nano}$ custom analysis software, (BMG Labtech GmbH, Germany). We performed Mann-Whitney test to compare the growth rate among conditions.

**Bacterial growth in solid HD**

Bacterial CFUs in HD were assessed in microtubes containing 400 µL of the diet of interest and 0.75–1 mm glass microbeads. Microtubes were inoculated with ~$10^4$ CFU of $Ap^{WJL}$ or $Lp^{NC8}$ or a ~$10^4$ CFU of a 1:1 mixture of $Ap^{WJL}$ and $Lp^{NC8}$ for coculture. To assess grow with larvae, 5 first-instar larvae, were added. The tubes were incubated at 25°C. After incubation, 600µL of PBS were added directly into the microtubes. Samples were homogenized with the Precellys 24 tissue homogenizer (Bertin Technologies, Montigny-le-Bretonneux, France). Lysates were diluted in PBS and plated on MRS. CFU counts were assessed as described above.

**Developmental timing determination**

Axenic adults were placed in sterile breeding cages overnight to lay eggs on sterile HD. The HD used to collect embryos always matched the experimental condition. Fresh axenic embryos were collected the next morning and seeded by pools of 40 in tubes containing 10mL of the HD to test. Unless otherwise stated, in mono-associated conditions a total of ~$10^5$ CFU of the strain of interest, washed on PBS, was inoculated on the substrate and the eggs. For bi-association ~$10^5$ CFU of a 1:1 mixture of $Ap^{WJL}$ and $Lp^{NC8}$ were inoculated. For heat killed (HK) conditions, washed cells of $Ap^{WJL}$ or $Lp^{NC8}$ were incubated 3h at 65°C. Once at room temperature, embryos were inoculated with ~$10^5$ or ~$10^9$ CFU. In the germ-free conditions, bacterial suspensions were replaced with sterile PBS. When testing the effect of bacterial by-products on developmental timing, 300 µL of supernatants of a 72h culture on complete HD of the strain of interest was added to the GF or mono-associated embryos. For the lactate supplementation experiments, DL-lactate, D-lactate or L-lactate (Sigma-Aldrich, Germany) were added to a final concentration of 0.6 g/L on GF or mono-associated eggs. For the amino acid cocktail supplementation experiment (Fig. 4), solid complete HD was supplemented with a solution containing the amino acid mixes described in Table S1.

After inoculation, the tubes were incubated at 25°C with 12/12-hour dark/light cycles. The emergence of pupae was scored every day until all pupae had emerged. The experiment was stopped when no pupae emerged after 30 days. Each gnotobiotic or nutritional condition was inoculated in five replicates. $D_{50}$ was determined using D50App (http://paulinejoncour.shinyapps.io/D50App) as described previously (Matos et al., 2017). $D_{50}$ heatmap represent the average of the five replicates of each gnotobiotic and nutritional condition. Fig 2K was done using the imagesc function on MATLAB (version 2016b. MathWorks, Natick, Massachusetts). Developmental timings are represented as boxplots showing the minimum, maximum and median where each point is a biological replicate. We performed Kruskal-Wallis test followed by uncorrected Dunn's tests to compare each gnotobiotic condition to GF or the condition indicated on the figure.

**Larval size measurements**

Axenic adults were placed in sterile breeding cages overnight to lay eggs on sterile HD. Fresh axenic embryos were collected the next morning and seeded by pools of 40 in tubes containing 10mL of complete HD. For the mono-associated conditions a total of ~$10^5$ CFU $Ap^{WJL}$ or $Lp^{NC8}$, washed on PBS, was inoculated on the substrate and the eggs. For biassociation ~$10^5$ CFU of a 1:1 mixture of $Ap^{WJL}$ and $Lp^{NC8}$ were inoculated. For the lactate supplementation experiments, DL-lactate was added to a final concentration of 0.6 g/L on $Ap^{WJL}$ mono-associated eggs. After inoculation, the tubes were incubated at 25°C with 12/12-hour dark/light cycles until collection of larvae. *Drosophila* larvae were randomly collected every day until day seven after inoculation and processed as described previously (Erkosar, 2015). Larval longitudinal length of individual larvae was quantified using ImageJ software.

**Microbial larval load in solid HD**

Axenic adults were placed in sterile breeding cages overnight to lay eggs on sterile HD. Fresh axenic embryos were collected the next morning and seeded by pools of 40 in tubes containing 10mL of complete HD supplemented with 0.08% of erioglaucine disodium salt (Sigma-Aldrich, Germany). For the mono-associated conditions a total of ~$10^5$ CFU $Ap^{WJL}$ or $Lp^{NC8}$, washed on PBS, were inoculated on the substrate and the eggs. For biassociation ~$10^5$ CFU of a 1:1 mixture of $Ap^{WJL}$ and $Lp^{NC8}$ were inoculated. When testing the effect of bacterial by-products on $Ap^{WJL}$ larval load, 300 µL of supernatants of a 72h culture on complete HD of the strain of interest was added to mono-associated embryos. After inoculation, the tubes were incubated at 25°C with 12/12-hour dark/light cycles until collection of larvae. *Drosophila* larvae were collected every day until five days after inoculation. We selected larvae with a blue gut to eliminate non-feeding individuals. Larvae were surface sterilize by rinsing once in ethanol 70% and twice in sterile PBS and placed in pools of 10 larvae/replicate/condition in 1.5 mL microtubes containing 500 µL of sterile PBS and 0.75–1 mm glass microbeads. Samples were homogenized with the Precellys 24 tissue homogenizer (Bertin Technologies, Montigny-le-Bretonneux, France). Lysates dilutions (in PBS) were plated on MRS and CFU counts were assessed as described above. Microbial larval loads are represented as dot plots where each point represents a biological replicate comprising the average microbial load of a pool of 10 larvae. We performed Mann-Whitney test to compare microbial loads in mono-association to microbial loads in biassociation for the strain of interest at each time point.

**DL-Lactate quantification**

Mono-cultures of $Ap^{WJL}$, $Lp^{NC8}$, $Lp^{WCFS1}$, $Lp^{WCFS1}\Delta$ldhDL, Af and co-cultures of $Ap^{WJL}$:$Lp^{NC8}$ and Af:$Lp^{NC8}$ were grown in liquid complete HD as described above. Samples were taken at time 0h and every 24h for 72 h. After centrifugation (5000 rpm, 5 min) to remove cells, D and L lactate concentrations were measured in the supernatants using the D-Lactate and L-Lactate Assay Kit, respectively (Megazyme, Pontcharra-sur-Turdine, France), following the manufacturers' recommendations.

**Amino acid quantification by HPLC**

In order to quantify Arg, Ile and Leu production in depleted media (Fig. 2H-J), PBS washed $Ap^{WJL}$, $Lp^{NC8}$ or $Ap^{WJL}$:$Lp^{NC8}$ were grown in liquid HDΔArg, HDΔIle or HDΔLeu as described above. Samples were collected every 24h for 72h. CFU counts were assessed as described above and supernatants were stored at -20°C until use.

To test total protein production by $Lp^{NC8}$ (Fig. S4E) PBS washed $Lp^{NC8}$ was grown in complete HD as described above. Supernatants were collected every 24h for 72h and stored at -20°C until use.

To test $Ap^{WJL}$ amino acid production upon DL-lactate supplementation (Fig. 4A-B), PBS washed $Ap^{WJL}$ was grown in complete HD supplemented or not with DL-lactate at final concentration of 20 g/L as described above. Supernatants were collected every 24h for 72h. CFU counts were assessed as described previously and supernatants were stored at -20°C until use.

Amino acid quantification was performed by HPLC from the supernatants. All proteinogenic amino acids were quantified except Cysteine, Tryptophan, Glutamine and Aspargine. Samples were crushed in 320 µl of ultra-pure water with a known quantity of norvaline used as the internal standard. Each sample was submitted to a classical protein hydrolysis in sealed glass tubes with Teflon-lined screw caps (6N HCl, 115°C, during 22h). After air vacuum removal, tubes were purged with nitrogen. All samples were stored at -20°C, and then mixed with 50 µL of ultra-pure water for amino acids analyses. Amino acid analysis was performed by HPLC (Agilent 1100; Agilent Technologies, Massy, France) with a guard cartridge and a reverse phase C18 column (Zorbax Eclipse-AAA 3.5 µm, 150 × 4.6 mm, Agilent Technologies). Prior to injection, the sample was buffered with borate at pH 10.2, and primary or secondary amino acids were derivatized with ortho-phthalaldehyde (OPA) or 9-fluorenylmethyl chloroformate (FMOC), respectively. The derivatization process, at room temperature, was automated using the Agilent 1313A autosampler. Separation was carried out at 40°C, with a flow rate of 2 mL/min, using 40 mM $NaH_2PO_4$ (eluent A, pH 7.8, adjusted with NaOH) as the polar phase and an acetonitrile/methanol/water mixture (45/45/10, v/v/v) as the non-polar phase (eluent B). A gradient was applied during chromatography, starting with 20% of B and increasing to 80% at the end. Detection was performed by a fluorescence detector set at 340 and 450 nm of excitation and emission wavelengths, respectively (266/305 nm for proline). These conditions do not allow for the detection and quantification of cysteine and tryptophan, so only 18 amino acids were quantified. For this quantification, norvaline was used as the internal standard and the response factor of each amino acid was determined using a 250 pmol/µl standard mix of amino acids. The software used was the ChemStation for LC 3D Systems (Agilent Technologies).

**Metabolite Profiling**

Samples were prepared from tubes inoculated as a DT experiment (see above) comprising 5 biological replicates per condition. Conditions included GF, Af and Af::Tn*ldh* (10B7) inoculated at ~$10^5$ CFU on complete HD in presence or not of a pool of 40 GF-eggs. For the lactate supplemented conditions, L-lactate (Sigma-Aldrich, Germany) was added to a final concentration of 0.6 g/L on mono-inoculated tubes (Fig. 6A). Tubes were incubated at 25°C with 12/12-hour dark/light cycles during 3 days. After incubation, a sample of minimum 100 mg was taken from the tubes. In the conditions including embryos, larvae were completely removed. Samples were stored at -80°C before sending to Metabolon Inc. (www.metabolon.com). Samples were extracted and prepared for analysis by Metabolon using standard solvent extraction method. The extracted samples were analysed using UltraHigh Performance Liquid Chromatography coupled to Tandem Mass Spectrometry. 321 compounds were identified by comparison to library entries of purified standards or recurrent unknown entities. Following log transformation and imputation of missing values, if any, with the minimum observed value for each compound, Welch's two-sample *t*-test was used to identify biochemicals that differed significantly between experimental groups.
